# Basolateral Amygdala Corticotrophin Releasing Factor Receptor 2 Interacts with Nonmuscle Myosin II to Destabilize Memory

**DOI:** 10.1101/2023.05.22.541732

**Authors:** Madalyn Hafenbreidel, Sherri B. Briggs, Meghana Arza, Shalakha Bonthu, Cadence Fisher, Annika Tiller, Alice B. Hall, Shayna Reed, Natasha Mayorga, Li Lin, Susan Khan, Michael D. Cameron, Gavin Rumbaugh, Courtney A. Miller

## Abstract

Inhibiting the actin motor ATPase nonmuscle myosin II (NMII) with blebbistatin (Blebb) in the basolateral amgydala (BLA) depolymerizes actin, resulting in an immediate, retrieval-independent disruption of methamphetamine (METH)-associated memory. The effect is highly selective, as NMII inhibition has no effect in other relevant brain regions (e.g. dorsal hippocampus [dPHC], nucleus accumbens [NAc]), nor does it interfere with associations for other aversive or appetitive stimuli, including cocaine (COC). To investigate a potential source of this specificity, pharmacokinetic differences in METH and COC brain exposure were examined. Replicating METH’s longer half-life with COC did not render the COC association susceptible to disruption by NMII inhibition. Therefore, transcriptional differences were next assessed. Comparative RNA-seq profiling in the BLA, dHPC and NAc following METH or COC conditioning identified *crhr2*, which encodes the corticotrophin releasing factor receptor 2 (CRF2), as uniquely upregulated by METH in the BLA. CRF2 antagonism with Astressin-2B (AS2B) had no effect on METH-associated memory after consolidation, allowing for determination of CRF2 influences on NMII-based susceptibility after METH conditioning. Pretreatment with AS2B occluded the ability of Blebb to disrupt an established METH-associated memory. Alternatively, the Blebb-induced, retrieval-independent memory disruption seen with METH was mimicked for COC when combined with CRF2 overexpression in the BLA and its ligand, UCN3 during conditioning. These results indicate that BLA CRF2 receptor activation during learning can prevent stabilization of the actin-myosin cytoskeleton supporting the memory, rendering it vulnerable to disruption via NMII inhibition. CRF2 represents an interesting target for BLA-dependent memory destabilization via downstream effects on NMII.

## Introduction

Substance use disorders (SUD) are long-lasting and perpetuated by associative memories, which can induce motivation to seek drugs. Memories are maintained, in part, by learning-induced structural changes to dendritic spines, which are mediated by actin dynamics [1]. We previously reported that actin remains uniquely dynamic in the basolateral amygdala (BLA) following methamphetamine (METH) treatment. Nonmuscle myosin II (NMII) is a molecular motor ATPase that drives actin polymerization to support this structural plasticity [2]. Inhibiting NMII with a single administration of blebbistatin (Blebb) arrests METH-induced actin dynamics and results in dendritic spine loss in the BLA, as well as an immediate, retrieval-independent disruption of established METH-associated memories and METH seeking that persists for at least one month [2–6]. The retrieval-independent effect of NMII inhibition is specific to METH and the BLA, as it does not interfere with memories for foot shock, food reward or other commonly abused drugs, including cocaine (COC) and does not disrupt METH memories when infused into dorsal hippocampus (dHPC) or nucleus accumbens [7,8]. We are currently developing a medication for METH use disorder based on these findings. However, the mechanisms underlying this highly specific effect are unknown.

The persistent susceptibility of a METH-associated memory to NMII inhibition was unexpected because actin-myosin dynamics are thought to rapidly stabilize after a learning event [1]. The sustained myosin-dependent actin dynamics in BLA spines [6] following METH treatment and the ability of Blebb to disrupt a METH-associated memory days to weeks after training [3,4] suggests that myosin remains uniquely active following METH exposure. To begin to approach the underlying mechanisms, we focused on the differential susceptibility of METH and COC to NMII inhibition. Given that METH and COC share a number of similarities [9,10], the specificity for METH is somewhat surprising. Very few comparative studies have investigated these illicit drugs, but one well-established difference is their pharmacokinetic properties. METH enters the brain at a higher concentration and has a longer half-life than COC [11–13]. We explored the hypothesis that longer brain exposure to METH outlasts the signaling cascade that typically inactivates NMII.

We further hypothesized that METH and COC are likely to induce a subset of transcripts unique from one another and between the BLA and other brain regions that support drug associations [14,15]. Identification of these differentially expressed genes (DEG) could provide insight into the selective vulnerability of METH-associated memories to NMII inhibition in the BLA. *Crhr2*, which encodes the corticotrophin releasing hormone or factor (CRF) receptor 2 (CRF2) was identified as uniquely upregulated in the BLA following METH treatment. CRF2 plays a key role in the brain’s stress response [16,17] and CRF, a ligand for both the CRF1 and CRF2 receptors, is released in the BLA following COC and METH administration [17,18]. In addition, CRF2 has been linked to NMII regulation outside the CNS [19]. Here we investigated the role of BLA CRF2 in METH and COC-associated learning and memory and its potential to drive differential susceptibility of the memories through NMII.

## Materials and Methods

### Subjects

Male C57BL/6 mice (Jackson Laboratory) weighing 25-30g were housed and handled as previously described [6,8]. Mice were housed four to a cage under a 12:12 light/dark cycle, with unlimited access to food and water, and were handled three days before behavioral testing. Sex as a biological variable was not examined as we previously found that NMII inhibition affects both sexes similarly [5]. Protocols were approved by the Institutional Animal Care and Use Committee at the Scripps Research Institute in accordance with National Institutes of Health guidelines.

### Drugs

Methamphetamine hydrochloride (METH: 2mg/kg, IP; Sigma-Aldrich) or cocaine hydrochloride (COC: 15mg/kg, IP; in the mini-pump, the concentration was 22.5mg/ml or 11.25mg/ml; NIDA) were dissolved in sterile 0.9% saline and administered at the designated doses. In some experiments, mice also received injections (IP) of vehicle (0.9% DMSO/25% hydropropyl β-cyclodextrin; HP βCD) or racemic (+/-) Blebbistatin (blebb; Tocris). Blebb was diluted to 1mg/ml in HP βCD vehicle, and administered at the dose of 10mg/kg, as previously described [4,6,8]. In some experiments, vehicle (sterile saline) or the selective CRF2 antagonist Astressin-2B [AS2B, Tocris; 20,21,22] was infused into BLA (1ug/0.5ul/side at a rate of 0.25ul/minute) 20 minutes before conditioning, or 15 minutes after the final conditioning session. AS2B dose, rate, and times of infusions were based on previous research [23–25]. Additionally, in other experiments, vehicle (sterile saline) or the selective CRF2 agonist Urocortin 3 (UCN3, Bachem) was infused into BLA (2ug/0.66ul/side at a rate of 0.3ul/minute) 15 minutes after the final conditioning session or 20 minutes before each PM conditioning session (CS+). UCN3 dose and rate were based on previous research [26–29].

### Conditioned place preference (CPP)

CPP was conducted similarly as previously described [3,30]. Briefly, mice were handled for three days before undergoing one or two 30 minute pretests, during which mice were allowed to freely explore all three CPP chambers. The final 15 minutes of the pretest were used to measure bias and to counterbalance groups for which chamber mice received drug (CS+) or saline (CS-). Mice did not have a significant preference for one chamber over the other (t’s<1.177, p’s>0.2513, r^2^<0.05787). However, mice that spent more than 90% of the total time in one chamber were excluded from analysis (n=4). Mice underwent conditioning over four consecutive days, with twice daily 30 minute conditioning sessions. To ensure there were no residual drug effects (Figure 1), saline was administered during the first (AM) and drug during the second (PM) conditioning session, which were separated by a minimum of four hours. In some experiments, mice were tested for memory retention during 15 minute free access sessions, similar to pretest, 48 hours after the final conditioning session, as previously done [3,8]. Mice were injected with saline (i.p.) before pre and post-tests, to mimic conditioning sessions procedures. All testing and conditioning were conducted in red or no light, and with a white noise generator (San Diego Instruments) set at ∼65-70dB near the CPP apparatus.

**Figure 1.**
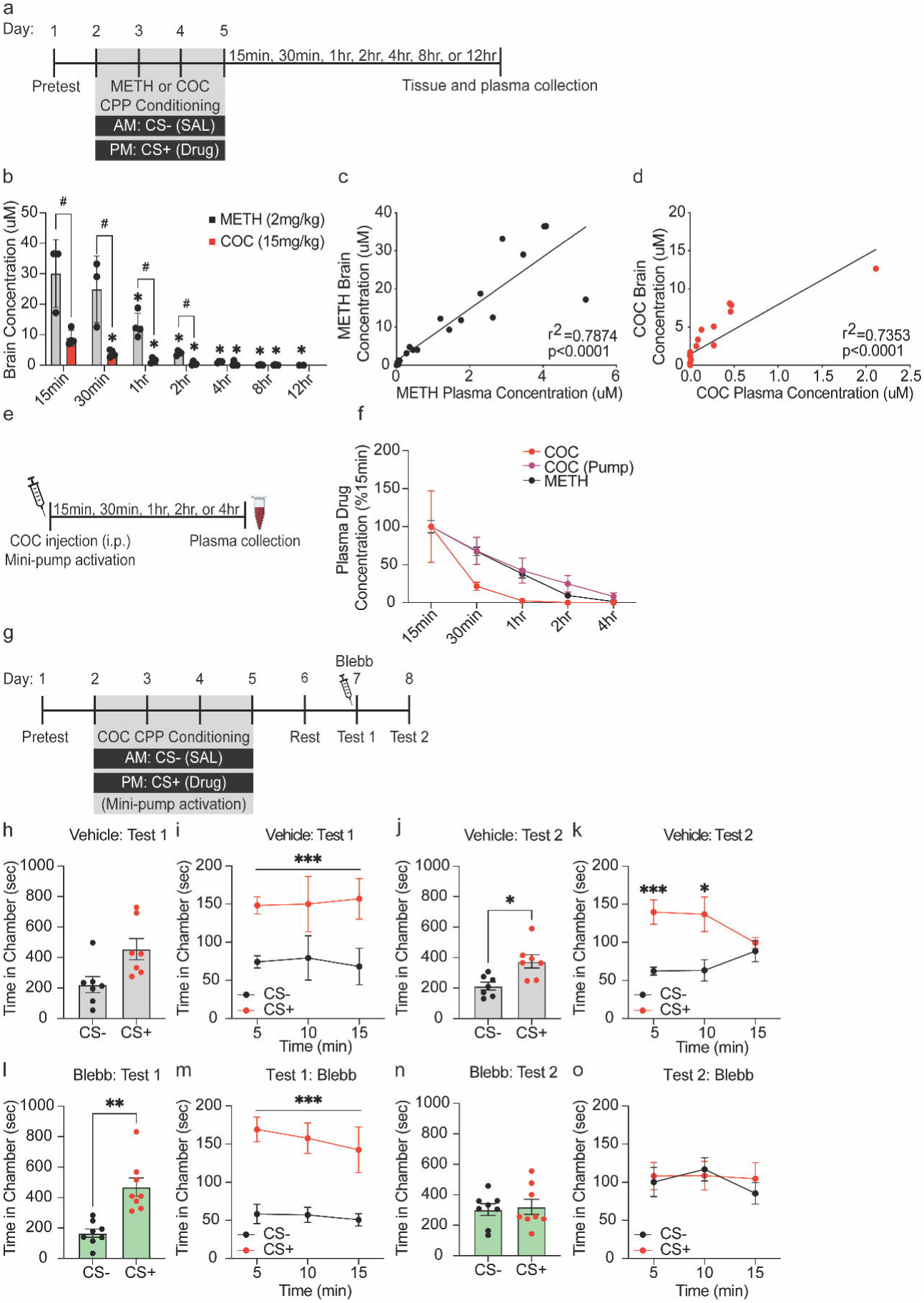
Susceptibility of METH, but not COC-associated memory to disruption by NMII inhibition is not due to half-life differences. a) Methods outline for b) METH and COC brain levels (n=3-5 for all time points, except 12 hr, where only METH samples were used, n=2;two-way repeated measures ANOVA revealed a significant effect of time (F_(6,35)_ =26.372, p<0.0001, η^2^=0.819), treatment ((F_(1,35)_ =64.809, p<0.0001, η^2^=0.649), and time by treatment interaction (F_(5,35)_ =10.709, p<0.0001, η^2^=0.605). A subsequent ANOVA confirmed that the concentration of COC was lower than METH and it was cleared more rapidly (15-minutes, F_(1,5)_ =14.381, p=0.013, η^2^=0.742; 30-minutes, F_(1,5)_ =15.570, p=0.011, η^2^=0.75; 1-hour, F_(1,6)_ =31.277, p=0.001, η^2^=0.839; 2 hours, F_(1,6)_ =59.906, p<0.0001, η^2^=0.909), but not 4- or 8-hours (F_(1,7)_ =4.074, p=0.083, η^2^=0.368; F_(1,5)_ =0.025, p=0.880, η^2^=0.005, respectively; METH, F_(6,16)_ =14.256, p<0.0001, η^2^=0.842; COC, F_(5,19)_ =32.307, p<0.0001, η^2^=0.895. *p<0.05 compared to 15-minutes, #p<0.05 between treatment groups). c) METH and d) COC brain concentrations are correlated with plasma levels (METH, r^2^=0.7874, n=23, p<0.0001, y=6.805x+1.189; COC, r^2^=0.7353, n=22, p<0.0001, y=6.506x+1.436). e) Methods outline for validating the programmable minipump, f) which mimics METH’s clearance rate with COC (n=2-4). ANOVA revealed a significant effect of time (F_(4,32)_ =13.869, p<0.0001, η^2^=0.634), but not treatment (F_(2,32)_ =2.625, p=0.088, η^2^=0.141), or time by treatment interaction (F_(8,32)_ =0.528, p=0.827, η^2^=0.117), indicating that plasma concentrations decreased as intended over time in COC Pump relative to METH IP. g) Methods outline for h-o. COC CPP in saline-treated mice the entire Test 1 (n=7; t_(6)_=2.021, p=0.0898, r^2^=0.4050) over h) and i) in 5-minute bins (ANOVA: significant effect of chamber (F_(1,36)_ =14.984, p<0.0001, η^2^=0.294), but not 5-minute time bin (F_(2,36)_ =0.011, p=0.989, η^2^=0.001) or chamber by bin interaction (F_(2,36)_ =0.076, p=0.927, η^2^=0.004), and Test 2 overall (j; t_(6)_=2.794, p=0.0314, r^2^=0.5653) and k) in 5-minute bins (ANOVA: significant effect of chamber (F_(1,36)_ =21.144, p<0.0001, η^2^=0.370) and a chamber by bin interaction (F_(2,36)_ =3.374, p=0.045, η^2^=0.158), but not bin (F_(2,36)_ =0.143, p=0.867, η^2^=0.008). This was confirmed by post hoc tests showing that time spent in the CS+ chamber is greater during the 5 (p<0.0001) and 10 (p=0.017), but not 15-minute (p=0.497) bins). COC CPP in Blebb-treated mice (n = 8) over l) the entire Test 1 (t_(7)_=3.714, p=0.0075, r^2^=0.6634) and m) in 5-minute bins (ANOVA: effect of chamber (F_(1,42)_ =48.726, p<0.001, η^2^=0.537), but not bin (F_(2,42)_ =0.479, p=0.623, η^2^=0.022) or a chamber by bin interaction (F_(2,42)_ =0.143, p=0.867, η^2^=0.007)), and test 2 n) overall (t_(7)_=0.2383, p=0.8285, r^2^=0.0080) and o) in 5=minute bins (ANOVA: no effect of bin (F_(2,42)_ =0.492, p=0.615, η^2^=0.023), chamber (F_(1,42)_ =0.174, p=0.678, η^2^=0.004) or chamber by bin interaction (F_(2,42)_ =0.174, p=0.678, η^2^=0.004). *p<0.05, **p<0.01, ***p<0.001. Error bars represent ± SEM.

### Mass spectrometry

Mice were anesthetized with isoflurane (Patterson Veterinary Supply Inc), and euthanized 15 minutes, 30 minutes, 1 hour, 2 hours, 4 hours, 8 hours, or 12 hours (post-injection) following the final METH conditioning session. Trunk blood was collected immediately into Eppendorf tubes that included 25ul of 10% EDTA (Ethylenediaminetetraacetic acid; to prevent coagulation) and stored on wet ice. Blood samples were later centrifuged for three minutes at 5,000 RPM to separate plasma from red blood cells. Plasma was collected into new tubes and stored at −80°C. Temporal lobe brain sections were simultaneously collected, and flash frozen with 2-methylbutane and stored at −80°C. METH or COC levels were quantified in plasma or brain samples by mass spectrometry using an ABSciex 5500 mass spectrometer using multiple reaction monitoring. Brain samples were homogenized in water and then immediately treated with 5-times (v:v) acetonitrile to extract the compound and precipitate cellular protein. Plasma samples were directly treated with acetonitrile. Samples were filtered through a 0.45µm filter plate prior to injection onto the LC-MS/MS.

### Experimental Manipulations

Full experimental details per each experiment can be found in the supplemental methods.

### Programable mini-pump implantation

Mice were implanted with programmable mini-pumps (i.p.; Primetech) programmed to deliver COC to mimic the clearance rate of METH. Mice were anesthetized with isoflurane (Kent Scientific) and mini-pumps were filled with COC, activated, and programmed (iPrecio). Pump tubing was inserted IP and sutured into place and the pump was implanted near the middle of the back. After surgery, mice were administered the analgesic Metacam (1.5mg/ml, 1-2 drops orally). Mice were allowed one week for recovery before CPP was conducted as described above. Specifically, mice received saline injections before the AM session, as usual, and then were injected with COC (IP; opposite side that the pump’s tube was inserted), and the pumps were programmed to begin expelling COC during the PM (CS+) sessions to mimic METH’s clearance curve (Figure 1). To verify the pumps were programmed correctly to mimic METH’s clearance rate, tail nick blood was collected 15 minutes, 30 minutes, 1 hour, 2 hours and 4 hours (post-injection) after the final conditioning session. Mice were warmed for 2-5 minutes under a heating lamp before being placed into a restraint tube. The tip of their tail was cut, or clot removed on subsequent collections, and blood was collected in a capillary plastic tube. Blood was centrifuged soon after and isolated plasma was stored at −80°C until concentrations were quantified by mass spectrometry, as described above. Once the pumps were validated, the experiment was replicated, but 48 hours after the final conditioning session, mice were injected with either vehicle or blebb (IP) 30 minutes before test 1. Mice were tested again 24 hours later in a drug-free state to assess memory retention.

### RNA Sequencing

To determine transcriptional differences between METH- and COC-associated learning, RNA-sequencing (RNAseq) was conducted on BLA, NAc, and dHPC collected following METH, COC, or saline (saline administered in AM and PM) CPP. Mice were euthanized 30 minutes after the final conditioning session to capture transcriptional changes during memory consolidation. Mice were anesthetized with isoflurane, after which brains were extracted, flash frozen with 2-methylbutane and stored at −80°C. Brain punches were later collected containing BLA, dHPC, or NAc. RNA was extracted using the miRVANA PARIS extraction kit (Life Technologies), as previously described [4,31]. RNA concentration was measured with a Qubit 3.0 Fluorometer and the Qubit RNA High Sensitivity Assay. RNAseq was conducted at the Scripps Florida Genomics Core. Total RNA quality was determined using the Agilent 2100 Bioanalyzer, and all RNA integrity Numbers (RINs) were ≥7.5. RNA libraries were prepared using the Illumina TruSeq RNA Library Preparation kit and protocol and sequenced on an Illumina NextSeq500 to yield at least ∼20million reads per sample. Initial analysis was conducted at the Scripps Florida Bioinformatics Core. Reads were mapped to the mouse genome (mouse-ENSEMBL-grcm38.r91:M.musculus-ENSEMBLE-GRCm38.r91) using the star version 2.5.2a aligner. Gene abundance was estimated with python version 2.7.11 and htseq version 0.11.0. Subsequent analysis identified genes that are significantly different (p<0.05) with at least +/-0.59 log2 (0.67>#>1.5) fold change between groups or structures. Genes that met this criterion then underwent additional analysis, including pathway analysis using Qiagen Ingenuity Pathway Analysis (IPA) software.

### Real time qualitative PCR (qPCR) and viral-mediated overexpression of CRF2

To determine if the optimal parameters for viral-mediated CRF2 expression at physiologically relevant levels (AAV5-mCrhr2-alpha) and to confirm the control virus (pAAV5- CMV-EGPF) did not alter CRF2 expression, 1:5, 1:10, and 1:30 dilutions were tested at 200 and 500nl volume injections in the BLA. Moreover, following the completion of the virus overexpression CPP experiments, mice were euthanized, and RNA expression was measured. RNA was extracted from BLA and hypothalamus (to measure probe efficacy and gross virus leakage) using the Zymo quick-RNA microprep kit. cDNA library was created from 100ng of total RNA using the TaqMan Fast Advanced Master Mix, Quantabio qScript cDNA SuperMix, and Taqman probes for crhr1 (mm00432670_m1) and crhr2 (mm00438305_m1). Data were normalized to the housekeeping Ubxn1 (mm00524986_m1; selected from RNAseq results showing it is not different between groups) using the ΔΔc_t_ method [32].

### Elevated Plus Maze and Open Field Test

To determine if intra-BLA infusions of AS2B following repeated METH injections attenuated anxiety-like behaviors, mice underwent testing in elevated plus maze (EPM) or open field (OF). Mice arrived, underwent cannula surgery, recovery, and handling similar to previous experiments. Next, mice received five daily injections of METH (IP and were placed into a clean, empty rat cage for 30 minutes to mimic CS+ conditioning). On the fourth and fifth days, 15 minutes after being removed from the empty cage (45 minutes after METH injection), mice were microinfused with vehicle, AS2B, or remained in their home cages. Mice then underwent EPM (day four) or OF (day five) testing 5-10 minutes after being microinfused. Testing was conducted as previously described [33] with some modifications. Briefly, during both EPM and OF testing, a white noise generator was used (San Diego Instruments; set to ∼70dB for apparatus) to mask external sounds and provide a constant noise level. Room and hallway were in red light. On day four, mice were placed in the center of the EPM (Med Associates) facing north toward an open arm, and activity was measured for five minutes using tracking software (EthoVision XT). On day five, mice were placed into the front right corner (facing center) of a custom-made open field box and activity was measured for five minutes using tracking software (EthoVision XT). Mice were then removed from the box and placed in their home cages. Boxes were cleaned with micro-90, and a clean novel object (mini-stapler) was placed in the center of the box to increase center interest in case of habituation during the first test. Mice were placed into the box again, in the same start location, for five minutes. Two OF tests were utilized to examine anxiety-like measures over ∼20 minutes to match the time period before mice underwent conditioning in experiment one.

### Corticosterone ELISA

Immediately following the end of the second OF test, mice were anesthetized with isoflurane and euthanized. Brains were collected for placement checks, and trunk blood was collected for corticosterone (CORT) ELISA analysis [using adapted methods from 34]. To include another time-point closer to when the mice were microinfused, an additional group of mice with previous CPP experience (from experiment five, remained in their home cages without disruption for ∼2.5 weeks to attenuate previous handing experience. Mice were then injected with METH (i.p.) and placed in a clean rat cage for 30 minutes, as done before. Forty-five minutes following the injection, mice received veh, AS2B or no infusion into BLA. Mice were anesthetized by isoflurane gas 20 minutes later, and trunk blood and brains were collected for CORT ELISA or placement checks, respectively. Trunk blood was collected into Eppendorf tubes with 100ul heparin (to prevent coagulation) and stored on wet ice until centrifuged for 10 minutes at 2500 RPM at 4°C to separate plasma from red blood cells. Plasma was collected into new tubes and stored at −80°C. CORT levels were determined by ELISA (Enzo Life Sciences) using the manufacturer’s protocol with an additional five minutes 98°C incubation in steroid dissociation reagent, which was previously found to be necessary to fully dissociate CORT from binding globulins [34].

### Data Analysis

Candidate RNAs were selected based on fold change (see above) and previous research. For behavioral experiments, a two-way repeated measures ANOVA was used to examine differences between time (bins within session) and chamber (CS+ or CS-), or a one-way ANOVA was used between treatment groups. A paired t-test was used to determine differences between chamber (CS+ vs CS-) during the posttest(s) and boxes (white vs black) during the pretest. All *post hoc* tests were conducted, where appropriate, using Tukey’s HSD test. Some mice were removed for having an initial bias (more than 90% time in one chamber during the pretest; n=4), using the ROUT method (Q=0.5%) to identify outliers or incorrect injector tip placement (n=7). Final group sizes are reported in figures. Effect size was determined using coefficient of determination (r^2^) for pair-wise comparisons or eta-squared (η^2^) for ANOVAs. After behavioral procedures, verification of injector tip location was performed on cresyl violet-stained or GFP expressing coronal sections.

## Results

### NMII susceptibility and the pharmacokinetics of METH and COC

We hypothesized that the longer brain exposure to METH may interfere with normal actin-myosin stabilization, resulting in the uniquely sustained susceptibility of METH-associated memory to NMII inhibition. To test this, we established the clearance rates of METH and COC, then used programmable mini-pumps to deliver COC at a rate that mimicked METH’s clearance and assessed the impact on COC-associated memory susceptibility. As repeated stimulant exposure can lead to a slowing of clearance [35], we determined the exact half-lives of METH and COC during CPP using doses that induce a reliable place preference with each drug [Figure 1a; 3,7]. As expected, METH levels were 3.4-fold higher than COC in the brain shortly after the final CPP conditioning session, despite the dose being 7.5-fold lower. Further, COC was cleared ∼4-fold faster (below detection at ∼8 hours for METH and ∼2 hours for COC; Figure 1b). Consistent with published results [11–13], COC had a half-life of 13.14-minutes and was cleared within ∼2 hours of administration, whereas METH had a half-life of 49.69-minutes and was cleared within ∼4 hours (Figure S1a-b). Brain levels correlated with plasma levels across the time course for both drugs (Figure 1c-d), enabling the use of plasma as a proxy for brain levels in subsequent experiments.

Using this information, we delivered COC through implantable, programmable mini-pumps at a rate that mimicked METH’s clearance. To confirm the accuracy, drug plasma levels were compared over time after an acute injection of cocaine COC (“COC IP”) and mini-pump infusion of COC (“COC Pump”; Figure 1e for methods; Figure S1c for plasma clearance rates). To validate the ability of COC Pump to mimic METH IP’s clearance rate, the differences in initial concentrations of METH and COC were accounted for by normalizing to their respective levels at 15-minutes post-administration (Figure 1f). Finally, the concentration of COC in the mini-pump was reduced by half (22.5mg/ml to 11.25mg/ml) to minimize the potential of inducing an aversion [36], however the clearance rate was maintained.

Next, mice were implanted with mini-pumps programmed as described above, with COC infused during CPP conditioning at the rate that mimicked METH’s clearance. Forty-eight hours after the last conditioning session, mice were injected with Blebb or vehicle (IP) and given a 15-minute memory retention test (Figure 1g). Vehicle-treated mice displayed a strong overall trend for a COC-associated memory during Test 1 (Figure 1h) with a significant preference for the COC-paired (CS+) chamber throughout the test session (Figure 1i); and a significant overall preference at Test 2 (Figure 1i). Analysis of within session performance during Test 2 suggests within session extinction occurred (Figure 1k). Blebb-treated mice displayed an overall preference during Test 1 (Figure 1l) and throughout the session (Figure 1m). However, Blebb-treated mice did not display a COC-associated memory during Test 2 (Figure 1n). These results indicate that mimicking METH’s clearance rate with COC did not render the COC-associated memory susceptible to immediate disruption by NMII inhibition. However, the results replicate our previous finding of disrupted COC-associated memory reconsolidation [7]. If Blebb treatment had resulted in rapid extinction, it would be expected that mice would display evidence of retrieval of the COC-associated memory during the initial part of the Test 2 session, which did not occur (Figure 1o). Overall, these results indicate that differing pharmacokinetics of METH and COC are unlikely to account for the unique susceptibility of METH-associated memory to NMII inhibition, but confirm our previous finding that NMII inhibition interferes with cocaine-associated memory reconsolidation [7].

### Differential gene expression is induced by METH and COC-associated learning

We next used RNA-seq to identify transcriptional profile differences accompanying METH- and COC-associated learning (Figure 2a). Profiles from the BLA were compared with two other brain regions that are also critical components of the neural circuit supporting drug-associated memory, but that do not share the susceptibility to METH-associated memory disruption with localized NMII inhibition, the NAc and dHPC [Figure 2b; 7]. RNA-seq was conducted on tissue collected 30-minutes after the last CPP conditioning session with METH, COC, or saline from BLA, NAc and dHPC. Bulk tissue was used to capture significant transcriptional changes across cell types, as the NMII-mediated effect on METH-associated memory has not been ascribed to a specific cell type. When comparing between brain regions within a given drug treatment, a similar number of genes were differentially expressed (Figure 2c i-iii). However, a striking difference emerged within the BLA when directly comparing differentially expressed genes (DEGs) between METH to COC (Figure 2c iv). Pathway analysis of DEGs in the BLA between METH and COC-treated mice identified a number of relevant functional and disease pathways, including psychological disorders, cell morphology and skeletal and muscular disorders, with the latter highlighting some myosin-related genes (*myl4, ppp1cb*, and *ppp1r1b*; Figure 2d-e). Fewer DEGs were identified when comparing METH or COC to saline control in the BLA (Figure 2c iv), indicating that METH and COC drive many transcriptional changes in opposing directions. Moreover, there were fewer DEGs in NAc or dHPC, as compared to BLA, between the METH and COC conditions (Figure 2c v-vi). The top 20 most highly expressed genes by treatment and brain region are shown in Figure S2a-i and the top ten DEGs for each comparison are presented in Figure S3a. Moreover, a number of genes were uniquely changed between brain regions (Figure 2f). There was less overlap in DEGs between brain regions for a given drug treatment comparison (Figure 2g).

**Figure 2.**
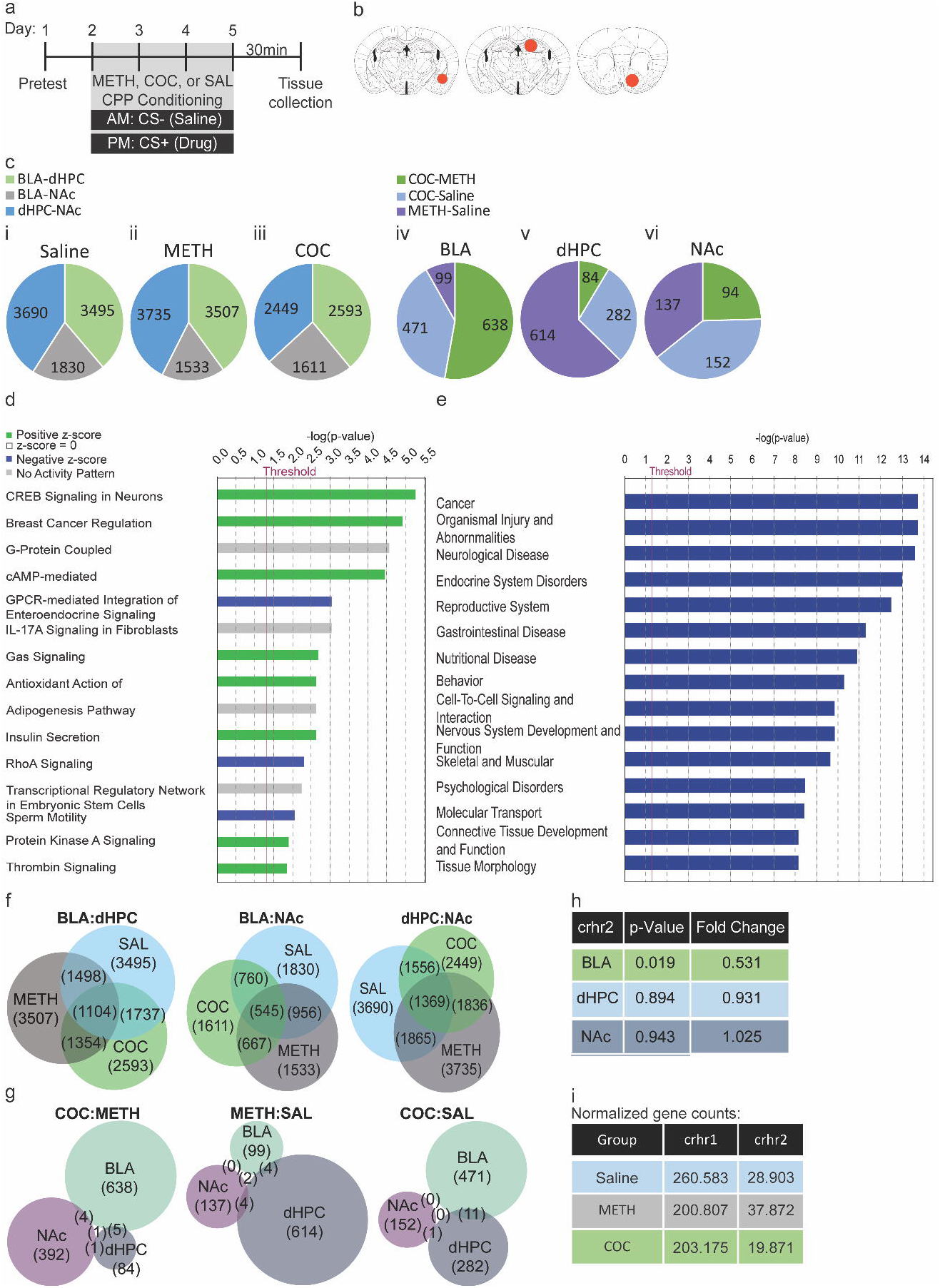
METH and COC induce unique expression of genes, such as *crhr2* (CRF2), in learning- and reward-related brain regions. a) Methods outline and b) representative regions of interest for RNAseq processing (n = 4-7). c) Differentially expressed genes by treatment (i-iii) and by brain region (iv-vi). d) function and e) disease pathways identified by Ingenuity Pathway Analysis (IPA). Uniquely changed genes when comparing f) brain regions and g) treatment. h) *crhr2* (CRF2) is significantly different in the BLA, but not dHPC or NAc, and i) is increased in METH compared to COC-treated mice.

### Role of BLA CRF2 in METH- and COC-associated learning

The transcriptional survey of DEGs (e.g., Figure S3b-c), identified *Crhr2*, which encodes Corticotrophin Releasing Factor (CRF) receptor 2 (CRF2), as selectively induced by METH compared to COC, in the BLA, but not NAc or dHPC (Figure 2h-i). IPA analysis also identified CRF signaling and its related network (Figure S4a) and signaling pathway (Figure S4b) at a DEG cutoff of p<0.05. CRF is important for drug-associated behaviors, as CRF1/2 antagonists attenuate stress-induced reinstatement following COC and heroin self-administration and disrupt COC CPP [37–40]. Further, a differential role has been reported for the CRF system for COC- and METH-associated behaviors [18,23,40,41]. Finally, CRF2 and its receptor-specific ligand, urocortin II, but not CRF1, have been linked to NMII signaling via effects on myosin light chain phosphorylation in the myometrium [19].

Before examining the potential influence of CRF2 on NMII, we assessed any potential role for CRF2 in METH-associated learning, as only locomotor sensitization had been previously tested [18,40,41]. Mice received intra-BLA infusions of vehicle or the CRF2-selective antagonist Astressin-2B (AS2B) 20-minutes before each of the four METH or COC conditioning sessions (Figure 3a and S5a). Consistent with previous reports [18,23,24,40,41], mice displayed COC-associated memory regardless of treatment (Figure 3b-c). Interestingly, a METH-associated memory was not present in mice that received intra-BLA Vehicle or AS2B infusions prior to each training session (Figure 3d-e). We repeated this experiment and found that again there was not a significant METH-associated memory in either group (Figure S6a-b), suggesting that formation of a METH-associated memory may be more sensitive to disruption by the process of intra-cranial infusion than a COC-associated memory. Together, these results indicate that CRF2 is not required for COC-associated learning but is unclear for METH-associated learning.

**Figure 3.**
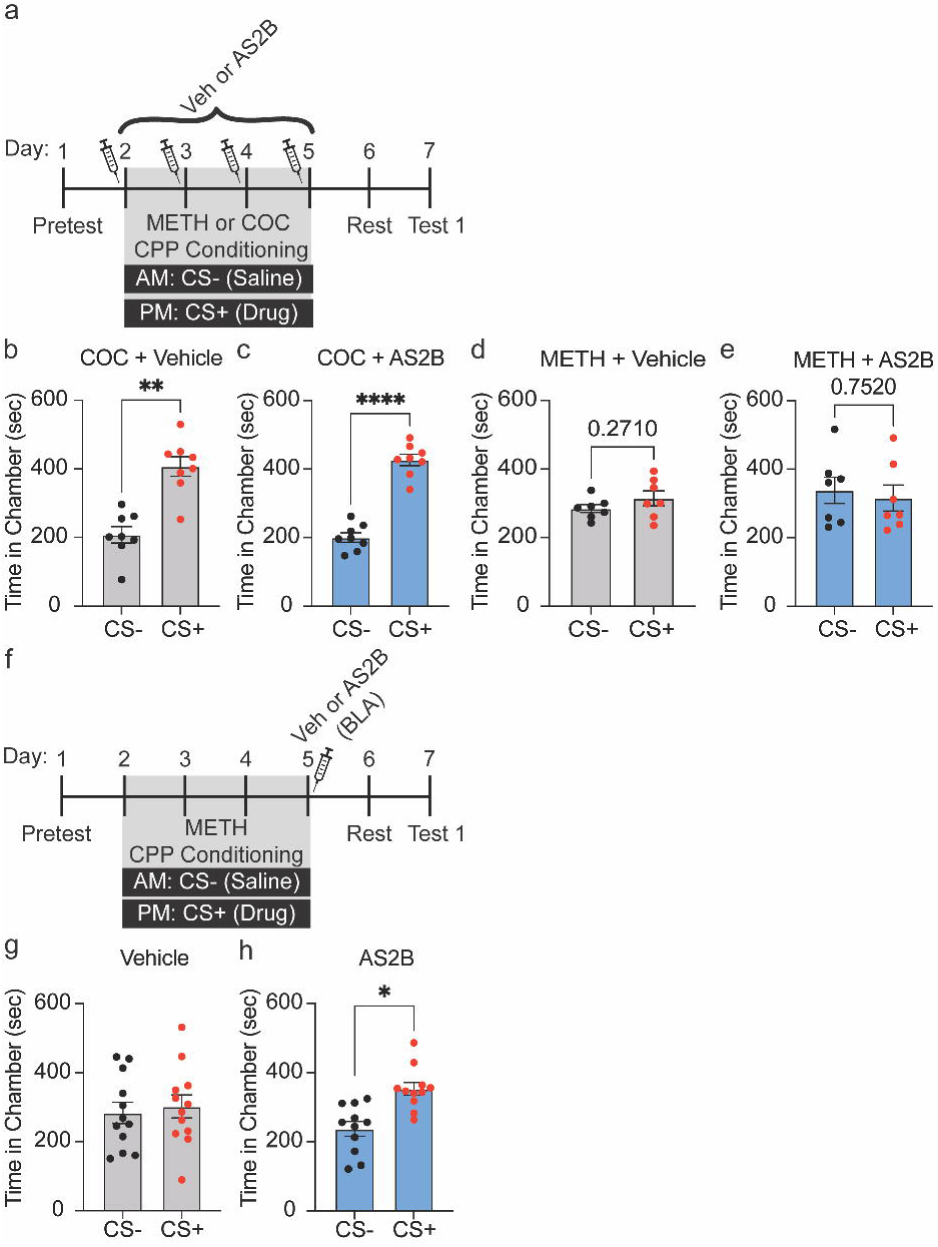
METH- but not COC-associated learning may involve BLA CRF2. a) methods outline. COC CPP following intra-BLA b) vehicle (n=8; t_(7)_=4.627, p=0.0024, r^2^=0.7536) or c) AS2B (n=8; t_(7)_=9.749, p<0.0001, r^2^=0.9314). METH CPP following intra-BLA d) vehicle (n=7; t_(6)_=1.212, p=0.2710, r^2^=0.1967) or e) AS2B (n=7; t_(6)_=0.3308, p=0.7520, r^2^=0.0179). f) methods outline. METH CPP following intra-BLA g) vehicle (n = 14; t_(11)_ =0.3145, p=0.7591, r^2^=0.0089) or h) AS2B (n=11; t_(10)_ =3.165, p=0.0101, r^2^=0.5004). *p<0.05, **p<0.01, ****p<0.0001. Error bars represent ± SEM.

### Intra-BLA CRF2 is not necessary for METH-associated learning once the memory is formed, but may have a role in anxiety-like behaviors

Our ultimate goal of testing for a potential role of CRF2 in NMII’s effects on an established METH-associated memory requires being able to inhibit CRF2 after the final conditioning session without interfering with expression of the METH association. Therefore, we next determined the impact, if any, of a single AS2B infusion 15-minutes after the final METH conditioning session on the subsequent expression of a METH-associated memory (Figure 3f, S5b). AS2B-treated, but not vehicle-treated mice expressed a METH-associated memory when tested 48 hours later (Figure 3g-h). These unexpected results suggested the potential of an anxiolytic effect of CRF2 inhibition in the BLA immediately following the final METH conditioning session. There is nothing in the literature to predict the likelihood of either possibility. Therefore, we performed elevated plus maze (EPM) and open field (OF) to assess the possibility of an anxiolytic effect but failed to find any remarkable outcomes of intra-BLA AS2B on anxiety-like behaviors in the context of prior METH exposure (supplemental results and supplemental figures S7-S9). It is possible that more sophisticated protocols or measures would reveal a dampening of anxiety-like behaviors with intra-BLA CRF2 antagonism.

### BLA CRF2 renders METH-associated memories selectively vulnerable to NMII inhibition

To mitigate the confounding effects of intra-BLA microinfusion, mice were habituated with mock infusions before each CPP conditioning session (Figure 4a, Figure S5c). Then, mice received intra-BLA infusions of vehicle or AS2B 15-minutes after the final conditioning session. Mock infusions were successful in protecting the formation of a METH-associated memory in vehicle-treated mice (Figure 4b). AS2B-treated mice also expressed a METH-associated memory (Figure 4c), enabling a subsequent test of the potential impact of CRF2 on NMII dynamics in support of METH-associated memory.

**Figure 4.**
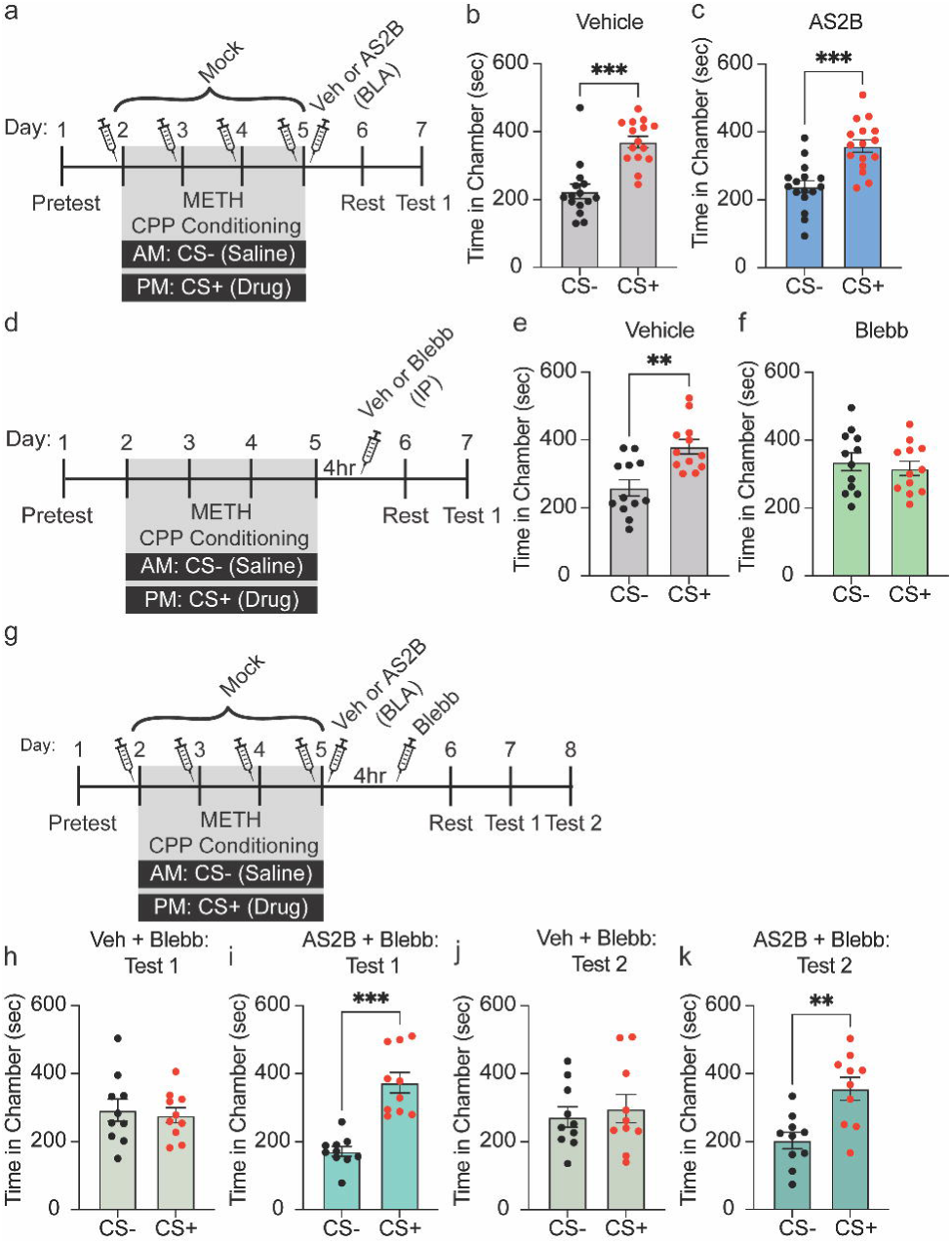
Intra-BLA CRF2 occludes the effect of systemic NMII inhibition on METH-associated memory. a) Methods outline for METH CPP following intra-BLA b) vehicle (n=15; t_(14)_ =4.058, p=0.0012, r^2^=0.5405) or c) AS2B after the last training session (n=16; t_(15)_ =3.859, p=0.0015, r^2^=0.4982). d) Methods outline for METH CPP following IP e) vehicle (n=12; t_(11)_ =3.184, p=0.0087, r^2^=0.4797) or f) Blebb 4 hours after the last training session (n=12; t_(11)_ =0.4608, p=0.6539, r^2^=0.0189). g) methods outline for METH CPP following h) vehicle + Blebb (n=10; t_(9)_=0.3334, p=0.7465, r^2^=0.0122) or i) AS2B + Blebb at Test 1 (n=10; t_(9)_=4.925, p=0.0008, r^2^=0.7293) and j-k) Test 2 (vehicle + Blebb: t_(9)_=0.3865, p=0.7081, r^2^=0.01632; AS2B + Blebb: t(_9_)=3.333, p=0.0088, r^2^=0.5524). **p<0.01, ***p<0.001. Error bars represent± SEM.

Prior experiments from our group have examined the impact of inhibiting NMII on established METH-associated memories anywhere from two days to two weeks after the final METH exposure [3]. Therefore, before the CRF2 and NMII interaction could be assessed, it was necessary to confirm that NMII inhibition is capable of disrupting METH-associated memories when delivered shortly after the final training session, in order to align with CRF2 timing. Blebb was administered four hours after the final METH conditioning session (Figure 4d). This time point was selected based on Figure 1’s data indicating that METH has cleared from the blood and brain by this point (Figure 1). When memory retention was tested 48 hours later, vehicle-treated mice displayed a significant METH-associated memory (Figure 4e), but Blebb-treated mice did not (Figure 4f). This established that the window of a METH-associated memory’s vulnerability to NMII inhibition stretches from ≤FOUR hours to ≥ two weeks post-training.

Taken together, these results enabled examination of the interaction between CRF2 and NMII, as we determined that CRF2 inhibition at this time point has no effect on METH-associated memory (Figure 4a-c), but NMII inhibition does (Figure 4f-g). We hypothesized that CRF2 antagonism (AS2B) would occlude the ability of NMII inhibition (Blebb) to disrupt METH-associated memory. To test this, mice again underwent mock infusions before each conditioning session for habituation, and then received intra-BLA infusions of vehicle or AS2B 15-minutes after the last conditioning session, followed by Blebb injection (IP) four hours later (Figure 4g, S5d). The METH-associated memory was disrupted in mice treated with the vehicle-Blebb combination (Figure 4h), replicating prior results [3]). However, METH-associated memory was protected in mice that were pre-treated with AS2B prior to Blebb (Figure 4i). Furthermore, this persisted 24 hours later during a second retention test (Figure 4j-k). Together, these results demonstrate the first potential mechanism underlying the BLA-dependent vulnerability of METH-associated memory to NMII inhibition.

### Activating the BLA CRF2 system renders COC-associated memory susceptible to NMII inhibition

Given our results demonstrating the interaction of CRF2 and NMII in supporting METH-associated memory, we next aimed to determine if a ‘gain-of-function’ approach could use the CRF2 system to render COC-associated memory susceptible to NMII inhibition. We first tested the potential to achieve this with CRF2 agonism in the BLA using urocortin 3 (UCN3), a CRF2-selective agonist. However, COC-associated memory remained intact when intra-BLA UCN3 was combined with Blebb (supplemental results and S10). These results suggest that a CRF2 receptor agonist alone following COC CPP conditioning is not sufficient to render the memory susceptible to disruption by NMII inhibition.

This result was, perhaps, not surprising because CRF2 is expressed at low levels throughout the brain, and particularly in the BLA [42]. Our RNAseq results indicate that COC reduced CRF2 expression even further (Figure 2i). Therefore, it is likely that too little receptor is present for UNC3 to act on. To address this, we overexpressed CRF2 in the BLA (AAV5-mCrhr2-alpha). The optimal titer and volume were first determined before behavioral testing (Supplemental Results and Figure S11-13). Next, a new cohort of mice received bilateral, intra-BLA injections of AAV5-Crhr2-alpha or control virus, followed by COC CPP training 30 days later and injection with vehicle or Blebb four hours after the final conditioning session (Figure S14a). There were no differences between groups: Figure S14b-e), indicating that simply overexpressing CRF2 in the BLA was insufficient to render the COC-associated memory susceptible to disruption. Additionally, we confirmed an increase of CRF2, but not CRF1 mRNA, in BLA with AAV5-mCrhr2-alpha (hypothalamus as control region; Figure S14f-g and Figure S15).

Finally, we tested the hypothesis that COC does not drive sufficient release of ligand in the BLA to activate the AAV5-mCrhr2-alpha-induced CRF2 receptors. Therefore, we included intra-BLA infusion of the CRF2-specific ligand, UCN3 (Figure 5a). Mice that received the combination of AAV5-mCrhr2-alpha/UNC3/Veh expressed a COC-associated memory during Test 1 (Figure 5b), but not in AAV5-mCrhr2-alpha/UNC3/Blebb-treated mice (Figure 5c). This persisted into Test 2, 24 hours later (Figure 5d-e). These results demonstrate that CRF2 overexpression in the BLA in combination with its selective ligand, UNC3, renders a COC-associated memory susceptible to disruption by NMII inhibition, mimicking the effect of NMII inhibition alone on a METH-associated memory. CRF2 mRNA was quantified to confirm AAV5-mCrhr2-alpha-mediated overexpression (Figure 5f-g; Figure 5h-i for representative images and placements).

**Figure 5.**
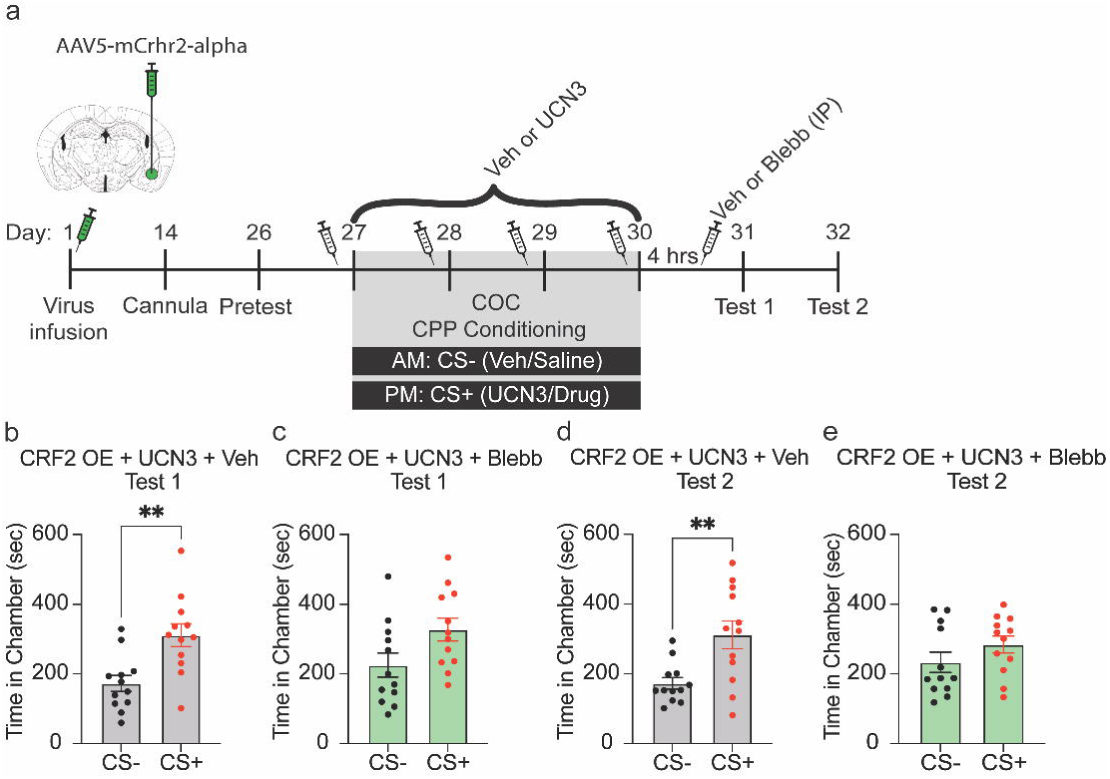
NMII inhibition plus viral overexpression and agonism of CRF2 in the BLA disrupts COC-associated memory. a) Methods outline. b) CRF2 overexpression (OE) + UCN3 + vehicle expressed a COC-associated memory (n=12; t_(11)_ =4.375, p=0.0011, r^2^=0.6351), c) but CRF2 OE + UCN3 + Blebb did not (n=12; t_(11)_ =1.775, p=0.1035, r^2^=0.2227). This persisted for two days (d-e; CRF2 OE + UCN3 + vehicle: t_(11)_ =3.926, p=0.0024, r^2^=0.5836 ; CRF2 OE + UCN3 + Blebb: t_(11)_ =1.243, p=0.2397, r^2^=0.1232). qPCR verified that CRF2 OE did result in an increase of CRF2 in the f) BLA (there was a significant difference in probe (F_(1,23)_ =23.109, p<0.0001, η^2^=0.501), treatment (F_(2,23)_ =3.879, p=0.035, η^2^=0.252) and a probe by treatment interaction (F_(2,23)_ =5.612, p=0.010, η^2^=0.0328). Subsequent ANOVAs determined that there was no difference in CRF1 (F_(2,12)_ =2.646, p=0.112, η^2^=0.306), but there was in CRF2 (F_(2,11)_ =4.428, p=0.039, η^2^=0.446). Post hoc analysis revealed that naïve mice are different than CRF2 overexpression + UCN3 + veh (p=0.031), but not CRF2 overexpression + UCN3 + Blebb (p=0.184), and CRF2 overexpression + UCN3 + Veh is not different than CRF2 overexpression + UCN3 + Blebb (p=0.389). Moreover, there was no difference between CRF1 and CRF2 in naïve mice (F_(1,4)_ =0.030, p=0.872, η^2^=0.007), but was in CRF2 overexpression + UCN3 + veh (F_(1,9)_ =20.064, p=0.002, η^2^=0.690) and CRF2 overexpression + UCN3 + Blebb (F_(1,10)_ =16.565, p=0.002, η^2^=0.624), but not g) hypothalamus (n=3-6; there was no significant effect of probe (F_(1,21)_ =0.320, p=0.577, η^2^=0.015) or probe by treatment interaction (F_(2,21)_ =0.135, p=0.874, η^2^=0.013) but there was an effect of treatment (F_(2,21)_ =4.475, p=0.024, η^2^=0.299). It is unclear why there is a treatment effect, but it is clear the CRF2 OE virus had no effect in this brain region and the probe was able to detect large increases in CRF2 in the BLA). h) Placement checks (CRF2 OE + UCN3 + vehicle = grey; CRF2 OE + UCN3 + Blebb = green). i) Representative images. **p<0.01. Error bars represent ± SEM.

## Discussion

Here we report that the difference in METH- and COC-associated memory susceptibility to disruption by NMII inhibition is not due to half-life. But, by examining transcriptional differences between METH- and COC-associated learning, we identified BLA CRF2 as a potential regulator. Interestingly, CRF1 is necessary for sensitization, CPP, and stress-induced reinstatement associated with COC, but not METH [18,23,24,40,41,43]. CRF2, on the other hand, is necessary for METH, but not COC, sensitization [18,40,41].

We found that CRF2 mRNA was selectively increased by METH, but not COC conditioning, in the BLA, but not dHPC or NAc. To determine the functional relevance, we examined the role of CRF2 in drug-associated learning, finding that it was not required for COC, but was unclear for METH. Regardless, CRF2 was not necessary once a METH-associated memory had formed. Moreover, inhibiting NMII just four hours after conditioning disrupted the METH-associated memory. When considered in the context of our prior findings [3,4], this indicates there is a persistent temporal window for METH-associated memory reliance on NMII, from the time of drug exposure until at least two weeks later, as opposed to a reliance induced later, during abstinence. These results allowed us to examine the interaction between CRF2 and NMII, finding that CRF2 is required in the BLA to render METH-associated memories selectively vulnerable to NMII inhibition in the absence of retrieval. In further support of the memory destabilizing effects of CRF2 on NMII, we made two discoveries. Intra-BLA administration of UCN3 in combination with NMII inhibition was insufficient to disrupt COC-associated memory. This was not surprising, as there is little CRF2 expression in the BLA under basal conditions [44] or following COC (Figure 2). However, administration of UCN3 during training, followed by NMII inhibition four hours after the last training session did disrupt a COC-associated memory with virus-driven expression of CRF2 in the BLA, confirming the ability of CRF2 to destabilize memory through NMII. These results identify the first upstream mechanism mediating our previously reported effects of NMII on METH-associated memories, an important mechanistic insight behind a novel therapeutic strategy being developed for METH use disorder [4,45,46]

Prior research on the role of BLA CRF2 effects on anxiety-like behaviors has been limited [37,47], perhaps because of the receptor’s very low basal expression levels. Depending on the task and species studied, CRF2 inhibition is either anxiolytic or benign [48,49]. Some of our findings suggested a possible link between BLA CRF2 and anxiety-associated effects. Mice that underwent COC conditioning expressed a strong place preference and AS2B had no effect on its expression, whereas mice that underwent METH CPP conditioning expressed a weaker association. The increase in BLA CRF2 expression associated with METH, but not COC CPP conditioning (Figure 3), may have induced a susceptibility to post-training microinfusion stress. However, our EPM and OF results suggest mice display some anxiety-like behaviors due to microinfusion, but there is no substantive effect of treatment. Perhaps more sensitive measurements would elicit differences. CRF2 is thought to have a bigger role in the post-stress response, or following the initial flight or fight response, and returning to homeostasis [47,50]. This is interesting considering post-traumatic stress disorder (PTSD) is often comorbid with SUDs and is a disease of disordered learning and HPA axis function. NMII inhibition does not have an immediate, retrieval-independent effect on fear-associated memory after consolidation [4], but it does disrupt fear memory reconsolidation [2]. As with COC-associated memory, fear memory destabilization and susceptibility to NM II inhibition could pot entially be achieved through manipulation of CRF2, opening a new therapeutic avenue.

Little is known regarding the mechanisms upstream of NMII that are responsible for the highly specific ability of NMII inhibition to drive retrieval-independent disruption of METH-associated memory. NMII is an activity-dependent molecular motor that drives synaptic plasticity through its time-limited effects on actin dynamics [2]. The sustained susceptibility of METH-associated memory to NMII inhibition longer after METH has cleared suggests that METH interferes with the normal inactivation of NMII. Indeed, we have shown that actin dynamics are sustained in BLA, but not dHPC, spines days after METH training [6]. The results presented here clearly implicate BLA CRF2 in the altered activity of NMII specifically associated with METH and the susceptibility of METH-associated memory to NMII inhibition (Figures 4). In further support of this, inducing the CRF2 system in the BLA was sufficient to induce susceptibility of a COC-associated memory to NMII inhibition (Figure 5).

Unfortunately, the CRF2 receptor has been far less studied than CRF1, particularly in the BLA, making it extremely difficult to predict how CRF2 may be producing a lasting influence on NMII function. However, unlike CRF1, CRF2 has clearly been linked to NMII. In human myometrial cells, UCN2 (another CRF2-selective ligand) triggers a signaling cascade that leads to NMII activation [19]. Each NMII molecule consists of two heavy chains, which bear the actin and ATP-binding sites, two essential light chains (ELC), which stabilize the myosin molecule, and two regulatory light chains (RLC) that, when phosphorylated, increase the actin-activated ATPase activity of NMII. UCN2 activates a signaling cascade that results in RLC phosphorylation [19]. This signaling cascade consists of proteins that are well-known to the field of synaptic plasticity, PKC, ERK1/2, RhoA and ROCK [51–54]. Further, they are all expressed in BLA and have been linked to METH [55–60].

How could CRF2 activation in the context of METH produce an effect on NMII function that is sustained and unique from COC? NMII is typically inactivated through dephosphorylation of the ELC by myosin light chain phosphatase (MLCP) [61]. Thus, sustained activation could be produced if the normal temporal window for recruitment of MLCP and inactivation of NMII is prevented by the long half-life of METH. However, mimicking the longer half-life of METH with COC was insufficient to induce susceptibility of a COC-associated memory to NMII inhibition (Figure 1). Another possibility is that CRF2 activation following METH exposure leads to the recruitment of a kinase that is not recruited by COC that phosphorylates (or drives some other activity modifying post-translational modification) at a unique site on NMII that results in sustained activation of the molecular motor. The result of such persistent NMII activation would be sustained susceptibility to Blebb, which targets the ATP-binding site, interfering with the energy needed for force generation to sustain actin dynamics. Identifying such a site will be a challenge, as a METH-induced modification could occur on any portion of the NMII protein. Further, given that such a modification would be driven by an exogenous agent not normally present in the brain (METH), it is possible that the enzyme capable of reversing the modification (e.g. a phosphatase) is simply not present in the BLA.

This study highlights the dramatic differences that two closely related psychostimulants, METH and COC, can have on the brain, with subregion-selectivity. This complexity is an important consideration, particularly given how common polysubstance use is. In addition, we established that the differences in susceptibility to disruption by NMII inhibition is not due to half-life differences, but a gene uniquely upregulated in the BLA following METH conditioning, CRF2, is required for the susceptibility of a METH-associated memory to NMII inhibition. Further, when CRF2 was overexpressed in the BLA and its ligand, UCN3, provided, a COC-associated memory became susceptible to NMII inhibition. These results strongly suggest that CRF2 contributes to altered function of NMII in the BLA in the context of METH. This provides new avenues to explore in the search for targets that selectively disrupt different pathogenic memories, including other SUDs and trauma-associated memories.

## Supporting information

Hafenbreidel et al., Supplemental methods and results

## Acknowledgments

We thank Dr. Stephanie E. Sillivan, Dr. Laszlo Radnai, Dr. Surya Pandey, Sarah Jamieson, Dr. Elizabeth M. Doncheck, Dr. Jennifer J. Tuscher, Pierce Herrmann, and Dr. John R. Mantsch for technical help. We also thank the Mouse Behavioral Core and Dr. Alicia Brantley for use of behavioral equipment, as well as the Scripps Genomics Core and Dr. Pabalu Karunadharma, and the Scripps Bioinformatics Core and Adrian Reich for their assistance with the RNAseq.

## Author Contributions

MH, SBB, and CAM designed the mini-pump experiments. MH and SBB conducted the behavior, with LL, SK and MDC performing the mass spectrometry. MH and CAM designed the RNAseq experiments, MH conducted the behavior and prepared the RNA. RNA was sequenced in the Scripps Genomics Core and analyzed by the Scripps Bioinformatics Core. MH and CAM analyzed the data further. MH and CAM designed the CRF2-related experiments, MH conducted the experiments. SR helped with the qPCR, and M H conducted them. MH, MA, SB, CF, AT, NM, and ABH did the associated histology and placements checks. GR helped with experimental design and virus design with help from SR. MH analyzed the data, and MH and CAM wrote and edited the manuscript. All authors reviewed the final content and approved it for publication.

## Funding

This research was supported by UH3 NS096833 and R01 DA049544 to CAM.

## Competing Interests

Authors have nothing to disclose.

## Notes

### Competing Interest Statement

The authors have declared no competing interest.

