## Supplementary material for "Basolateral Amygdala Corticotrophin Releasing Factor Receptor 2 Interacts with Nonmuscle Myosin II to Destabilize Memory": Hafenbreidel et al., Supplemental methods and results

**Experimental Manipulations**

*Experiment one.* To determine the role of intra-BLA CRF2 in METH- or COC-associated memories, CRF2 was inhibited prior to every CPP conditioning session. Following one week of acclamation after arrival, mice were implanted with two single-barrel 26G guide cannula aimed bilaterally at the BLA (AP, -1.5mm, ML, +/-3.2mm, DV -3.7mm from skull) as previously reported [1] using the neurostar robotic stereotaxic equipment. Metacam, an analgesic, was administered after surgery. Mice were given one week for recovery before being handled for three days, which included habituation to microinfusion procedures. Mice were handled as previously described [2] before being transported to the room in which infusions would be received. On day one, mice were scruffed and dust caps were removed; on day two, injectors extending 1mm past the guide cannula were lowered to the infusion site for 1-2 minutes; and on day three, saline was infused at the same rate and volume as during drug manipulation to allow mice to adapt to changes in cranial pressure and mechanical stimulation. The following day, mice underwent a pretest, as described above, except mice had infusers lowered 20 minutes before testing. During the next four days, mice were conditioned as described above except mice received microinfusions of vehicle or AS2B 20 minutes prior to every conditioning session (CS- in AM and CS+ in PM, METH or COC). Memory retention was tested 48 hours later.

*Experiment two.* To determine if intra-BLA CRF2 is necessary for memory acquisition or after the memory was formed, CRF2 were inhibited only after the final conditioning session. Experimental procedures were identical to experiment one except mice were only infused 15 minutes after the final CS+ conditioning session. Memory retention was tested 48 hours later.

*Experiment three.* To control for the potential stressful effect on memory expression of a single microinfusion into BLA 15 minutes after the final conditioning session (when mice are still under the influence of METH), experiment two was replicated except mice received mock infusions 20 minutes prior to each conditioning session, similar to experiment one. A mock infusion was conducted identically as a normal infusion, but no liquid was expelled from the infusers. Mice were infused with vehicle or AS2B 15 minutes after the final conditioning session. Memory retention was tested 48 hours later.

*Experiment four.* Previously, we found that NMII inhibition can disrupt METH-associated memories when injected 30 minutes or 24 hours before testing [1,3,4]. However, it was unclear whether memory retention would be disrupted during the consolidation window following the final day of conditioning. Mice underwent METH CPP as described but were injected with vehicle or blebb (IP) four hours after the final conditioning session. Memory retention was tested 48 hours later.

*Experiment five:* To determine the interaction between CRF2 and NMII, experiment three was replicated, but mice also received blebb injections (IP) four hours after the final conditioning session (from the time of METH injection). Moreover, to determine if CRF2 is bidirectionally mediating drug-associated learning, experiment five was replicated except mice were conditioned with COC and infused with vehicle or the CRF2 selective agonist Urocortin 3 (UCN3) into BLA 15 minutes after the final conditioning session, then four hours later (from time of METH injection) mice received an injection of blebb (IP).

Experiment six: The CRF2 overexpression virus **(**(pAAV [exp]-CMV.mCrhr2[NM_001288619.1]:RES:NLS-EGFP; AAV5-mCrhr2-alpha) was designed in house through vector builder. The control virus (pAAV.CMV.PI.EGFP.WPRE.bGH; pAAV5-CMV-EGFP) was purchased through Addgene. Following one week of acclamation after arrival, mice were injected with virus aimed bilaterally at the BLA (AP, -1.6mm, ML, +/-3.45mm, DV -5.05mm from skull). Surgery was conducted similar to cannula surgeries except no headcaps were used.

Mice were given 30 days for virus expression. Next, the experiment was conducted similar to experiment five. Mice were injected with blebb or vehicle (IP) four hours after the final conditioning session. Two additional controls were included that were also implanted with cannula and received saline microinfusions in the AM session (saline IP injections) and UCN3 injections in the afternoon (COC IP injections), and then four hours after the final conditioning session were injected with blebb or vehicle (IP). UCN3 was administered during this time frame to attempt to mimic METH's course of action following injection.

**Supplemental results**

**NMII Susceptibility and the pharmacokinetics of METH and COC**

Using the determined clearance rates, we preformed modeling to determine a dosing rate of COC that would mimic METH's clearance with implantable, programmable minipumps for drug delivery. To confirm accurate modeling of METH's clearance rate with COC, drug plasma levels were compared over time (15 minutes to 4 hours) after an acute injection of cocaine COC ("COC IP") and mini-pump infusion of COC ("COC Pump", Figure 1e). Two-way repeated measures ANOVA confirmed this, revealing significant effect of time ($F_{(4,32)}$=50.510, p<0.0001, $\eta^2$=0.863), treatment (($F_{(2,32)}$=100.208, p<0.0001, $\eta^2$=0.862), and time by treatment interaction ($F_{(8,32)}$=10.709, p<0.0001, $\eta^2$=0.802; Figure S1c). To confirm there were differences between treatment groups, a subsequent ANOVA revealed a difference between groups at all time points except 4 hours (15 minutes, $F_{(2,6)}$=23.964, p=0.001, $\eta^2$=0.889; 30 minutes, $F_{(2,6)}$=77.084, p<0.0001, $\eta^2$=0.963; 1 hour, $F_{(2,7)}$=24.521, p=0.001, $\eta^2$=0.875; 2 hours, $F_{(2,7)}$=13.349, p=0.004, $\eta^2$=0.792; 4 hours, $F_{(2,6)}$=4.560, p=0.062, $\eta^2$=0.603). Post hoc tests confirmed that METH IP has a higher concentration than COC IP at all time points (p's<0.004) and COC pump (p's<0.008,) except at 2 hours (p=0.913), and COC IP was not different from COC pump at 15 minutes (p=0.581). These differences were intentional, as COC becomes aversive at higher concentrations [5]. At 30 minutes and 2 hours, COC Pump was higher than COC IP (p's=0.042 and 0.022, respectively), confirming the programmed pump maintains COC at appropriate, detectable levels for the same time course as METH.

**Intra-BLA CRF2 does not impact EPM and OF**

To explore the possibility of an anxiolytic effect of CRF2 inhibition in the context of METH exposure, a protocol similar to the CPP injection regimen was used in which mice were placed in a novel environment for 30 minutes after a METH injection. This was repeated for 5 days (Figure S7a for cannula placements, Figure S7b for methods timeline). Fifteen minutes after removal from the novel environment on the fourth day, mice were given intra-BLA infusions of vehicle, AS2B, or no infusion (remained in homecage). Anxiety-like behavior was then examined in EPM 5 minutes later. These methods were repeated the following day with OF. Mice were given two 5 minute sessions in OF (~6 minutes between tests for cleaning), after which plasma was collected for corticosterone concentration analysis (~25 minutes after infusions). Another cohort of mice underwent similar methods, but were euthanized 15 minutes following vehicle, AS2B, or no infusions without any behavioral testing. Corticosterone concentrations were unchanged (Figure S7c, ~25 minutes post: $F_{(2,28)}=0.3795$, $p=0.6877$, $\eta^2=0.0648$; Figure S7d, ~15 minutes post: $F_{(2,19)}=0.9926$, $p=0.3980$, $\eta^2=0.0946$).

In EPM, there was a difference in latency to the open arms (Figure S7e, $F_{(2,21)}=3.709$, $p=0.0417$, $\eta^2=0.2610$), but post-hoc tests failed to identify a difference between groups. There were no differences in latency to closed arms (Figure S7f, $F_{(2,29)}=0.2940$, $p=0.7475$, $\eta^2=0.01987$). Total distance moved was not different between groups (Figure S7g, $F_{(2,29)}=0.0630$, $p=0.9391$, $\eta^2=0.0043$), indicating no effect on locomotion as expected as all groups received METH injections. Finally, frequency (Figure S7h, open: $F_{(2,29)}=0.0304$, $p=0.9701$, $\eta^2=0.00209$; Figure S7i, closed: $F_{(2,29)}=0.1209$, $p=0.8866$, $\eta^2=0.0083$) and duration

(Figure S7j, open: $F_{(2,29)}$=0.1407, p=0.8693, $\eta^2$=0.0096; Figure S7k, closed: $F_{(2,29)}$=0.5814, p=0.5655, $\eta^2$=0.0386) to enter arms was not different between groups.

In OF Test 1, center frequency and duration were not different between groups (Figure S8a, frequency: $F_{(2,28)}$=0.0473, p=0.9539, $\eta^2$=0.0034; Figure S8b, duration: $F_{(2,27)}$=0.2837, p=0.7552, $\eta^2$=0.0206), but wall frequency and duration were different (Figure S8c, frequency: $F_{(2,28)}$=4.288, p=0.0237, $\eta^2$=0.2345; Figure S8d, duration: $F_{(2,28)}$=5.132, p=0.0126, $\eta^2$=0.2683). No infusion-treated mice had higher wall frequency (p=0.0246) and duration (p=0.0104) compared to vehicle-treated mice, but AS2B-treated mice were not different than either group (p's>0.05). Latency to center was not different between groups (Figure S8e, $F_{(2,28)}$=2.396, p=0.1095, $\eta^2$=0.1462) nor latency to the wall (Figure S8f, $F_{(2,27)}$=0.3173, p=0.7308, $\eta^2$=0.02297). Distance and Velocity were not different between groups, indicating no effect on locomotion (Figure S8g, Distance: $F_{(2,28)}$=1.578, p=0.2241, $\eta^2$=0.1013; Figure S8h, Velocity: $F_{(2,28)}$=1.309, p=0.2860, $\eta^2$=0.0855). However, when within session is analyzed to examine the first minute of the session in order to assess initial effects on stress-like behaviors, there are differences in treatment groups in distance moved (Figure S8i, distance, $F_{(2,28)}$=4.097, p=0.0275, $\eta^2$=0.2264). Post hoc tests confirm that infusions have an effect as no infusion-treated mice moved less than vehicle-treated mice (p=0.0426) but were not different than AS2B-treated mice (p=0.9816), and there is a nonsignificant trend for vehicle-treated mice to locomote more than AS2B-treated mice (p=0.0638). Similarly there were group differences in velocity during the first minute, (Figure S8j, velocity, $F_{(2,28)}$=4.530, p=0.0198, $\eta^2$=0.2445). Post hoc tests confirm that infusions have an effect as no infusion-treated mice moved less than vehicle-treated mice (p=0.0184) but were not different than AS2B-treated mice (p=0.6779), and vehicle-treated mice were the same as AS2B-treated mice (p=0.1193). Together these results suggest that during test 1 vehicle-treated mice are locomoting more and spending less time and frequency near the wall than no-infusion mice, and AS2B infusions might have partly reversed these changes. Overall, these results suggest that microinfusion decrease anxiety-like behaviors.

During the second open field test, which is closer to when mice would undergo conditioning during previous experiments timewise, also included a novel object (mini-stapler) in the middle of the box to increase interest if habitation occurred during the first test. Center frequency was not different between groups (Figure S9a, $F_{(2,28)}=0.3072$, $p=0.7380$, $\eta^2=0.0215$) or duration (Figure S9b, $F_{(2,28)}=0.8026$, $p=0.4582$, $\eta^2=0.0542$). Wall frequency was also not different between groups (Figure S9c, $F_{(2,28)}=2.616$, $p=0.0909$, $\eta^2=0.1574$), but duration was (Figure S9d, $F_{(2,27)}=6.127$, $p=0.0064$, $\eta^2=0.3122$). Post hoc tests confirmed vehicle-treated mice spent less time by the walls than no infusion-treated mice ($p=0.0440$) and AS2B-treated mice ($p=0.0064$), but AS2B- and no infusion-treated mice were not different (0.6951) suggesting intra-BLA blockade of CRF2 can reverse the anxiolytic effect of a vehicle infusion. Frequency and duration around the novel object (middle) were not different between groups (Figure S9e, frequency: $F_{(2,28)}=0.0943$, $p=0.9103$, $\eta^2=0.0067$; Figure S9f, duration: $F_{(2,27)}=0.8160$, $p=0.4528$, $\eta^2=0.0570$). Latency to center (Figure S9g, $F_{(2,26)}=0.1825$, $p=0.8343$, $\eta^2=0.01384$), wall (Figure S9h, $F_{(2,28)}=2.154$, $p=0.1349$, $\eta^2=0.1333$), or middle (Figure S9i, $F_{(2,27)}=0.8935$, $p=0.4210$, $\eta^2=0.06208$) were not different between groups. Distance moved was not different between groups (Figure S9j, $F_{(2,28)}=0.1189$, $p=0.8884$, $\eta^2=0.0084$) nor was velocity (Figure S9k; $F_{(2,28)}=0.1172$, $p=0.8899$, $\eta^2=0.0083$). However, like test 1, when the first minute of the session is analyzed, differences between groups are observed (Figure S9l, $F_{(2,27)}=5.563$, $p=0.0095$, $\eta^2=0.2918$). Post hoc tests reveal that no Infusion-treated mice locomoted less than vehicle- ($p=0.0396$) and AS2B-treated mice ($p=0.0103$), but vehicle- and AS2B-treated mice were not different ($p=0.7825$) suggesting microinfusions affects initial locomotion but blocking CRF2 has no effect. Similarly, with velocity, there was a difference between groups when the first minute was analyzed (Figure S9m, $F_{(2,27)}=5.620$, $p=0.0091$, $\eta^2=0.2940$) and had a similar pattern of results as distance: No infusion-treated mice had lower velocity than vehicle- ($p=0.0409$) and AS2B-treated mice ($p=0.0096$), but vehicle- and AS2B-treated mice were not different ($p=0.7559$).

Here, we found that infusing vehicle in the BLA had a slight anxiolytic effect, which was slightly reversed in some measures by inhibiting CRF2. Mice were habituated to microinfusion procedures before the start of the experiment, and mice were infused before the EPM which was the day before, which could explain why vehicle microinfusion treatment resulted in a more anxiolytic effect compared to an expected axiogenetic effect. This could explain too why habituation before each conditioning session (e.g., Figure 4) was able to rescue the METH-associated memory. EMP and OF were not counterbalanced because of technical limitations with collecting the plasma for the corticosterone measurement. Moreover, previous research has suggested that CRF2's role in anxiety-like behaviors was mixed depending on measurement, brain region and species used, from having no effect to being anxiolytic [6,7]. Our results suggest CRF2 blockade in the BLA did little to anxiety-like behaviors as measured here.

**Activating the BLA CRF2 system renders COC-associated memory susceptible to NMII inhibition**

Mice received mock infusions prior to each COC conditioning session (Figure S10a for cannula placements, Figure S10b for methods timeline), then intra-BLA infusions of vehicle or CRF2 agonist UCN3 15 minutes after the final conditioning session and blebb (IP) four hours later. All groups expressed a METH-associated memory 48 hours later during a memory retention test (Figure S10c, vehicle + vehicle, $t_{(13)}=3.047$, $p=0.0094$, $r^2=0.4167$; Figure S10d, vehicle + blebb, $t_{(13)}=2.930$, $p=0.0117$, $r^2=0.3978$; Figure S10e, UCN3 + vehicle, $t_{(13)}=3.512$, $p=0.0038$, $r^2=0.4869$; Figure S10f, UCN3 + blebb, $t_{(12)}=3.375$, $p=0.0055$, $r^2=0.4970$) indicating that potentiating CRF2 plus inhibiting NMII had no effect on COC-associated memories.

To determine the optimal dilution of the virus used to overexpress (OE) CRF2, mice were infused with 1:30, 1:10, or 1:5 dilutions of AAV5-mCrhr2-alpha bilaterally. Additional groups of 1:10 dilution were included to examine the spread of virus which included mice infused with 200 or 500nl of AAV5-mCrhr2-alphaα or control (pAAV5-CMV-EGFP) virus (Figure S11a). Thirty days later, mice were tested in OF and then euthanized for placement checks and qPCR verification. In the BLA (Figure S11b), qPCR verified that CRF2 was significantly increased (Probe: $F_{(1,32)}=9.085$, $p=0.005$, $\eta^2=0.221$; Dilution: $F_{(3,32)}=6.088$, $p=0.002$, $\eta^2=0.363$; probe by dilution interaction: $F_{(3,32)}=6.164$, $p=0.002$, $\eta^2=0.366$) compared to CRF1 at the 1:5 dilution (Naïve: $F_{(1,8)}<0.0001$, $p=0.994$, $\eta^2<0.0001$; 1:30: $F_{(1,8)}=1.300$, $p=0.287$, $\eta^2=0.140$; 1:10: $F_{(1,8)}=3.776$, $p=0.088$, $\eta^2=0.321$; 1:5: $F_{(1,8)}=7.023$, $p=0.029$, $\eta^2=0.467$). A control brain region was included, and CRF2 and CRF1 expression was not different between any dilution in the hypothalamus (Figure S11c; Probe: $F_{(1,29)}=0.977$, $p=0.331$, $\eta^2=0.033$; Dilution: $F_{(3,29)}=1.032$, $p=0.393$, $\eta^2=0.096$; probe by dilution interaction: $F_{(3,29)}=0.617$, $p=0.609$, $\eta^2=0.060$). Representative images are in Figure S11d. From these results we concluded that the optimal dilution was 1:10 as we did not want to induce extra-physiological levels of CRF2 in the BLA, while still maintaining good virus expression.

Next, we determined the optimal infusion volume to restrict OE to the BLA, testing 200 and 500nl. Thirty days after virus injection, mice were tested in OF and then euthanized for placement checks and qPCR verification. In the BLA (Figure S12ad), there was a significant effect of probe ($F_{(1,12)}=25.584$, $p<0.0001$, $\eta^2=0.681$), volume ($F_{(1,12)}=8.986$, $p<0.0001$, $\eta^2=0.428$), and probe by volume interaction ($F_{(1,12)}=7.436$, $p=0.018$, $\eta^2=0.383$). Subsequent ANOVAs reveal there are no differences between CRF1 expression ($F_{(1,6)}=2.135$, $p=0.194$, $\eta^2=0.262$), but a significant difference between CRF2 expression ($F_{(1,6)}=8.264$, $p=0.028$, $\eta^2=0.579$). Moreover, there was a difference between CRF2 OE virus 200nl between CRF1 and CRF2 ($F_{(1,6)}=11.033$, $p=0.016$, $\eta^2=0.648$) and 500nl ($F_{(1,6)}=17.279$, $p=0.006$, $\eta^2=0.742$). Overall, these results confirm that CRF2 was overexpressed by the CRF2 OE virus in the BLA. We also examined the hypothalamus (Figure S12b). There was no significant differences by probe ($F_{(1,12)}$=3.323, p=0.0.93, $\eta^2$=0.217), infusion volume ($F_{(1,12)}$=1.552, p=0.237, $\eta^2$=0.115), or probe by infusion volume interaction ($F_{(1,12)}$=0.782, p=0.394, $\eta^2$=0.061). Based on the representative images (Figure S12c) and qPCR, we continued our experiments with 1:10 dilution to limit the virus overexpression to keep it from super physiological levels, and 500nl in order to get the best spread within the BLA.

Additionally, we examined behavior in OF with different dilutions and infusion volumes of AAV5-mCrhr2-alpha. There were no significant differences between groups when distance traveled was examined (Figure S13a; $F_{(2,23)}$=0.3625, p=0.6999, $\eta^2$=0.03056) or velocity (Figure S13b; $F_{(2,23)}$=0.2916, p=0.7298, $\eta^2$=0.02473). Finally when the first minute was examined to look at initial anxiety, there were no differences in distance traveled (Figure S13c; $F_{(2,23)}$=0.1019, p=0.9035, $\eta^2$=0.0088) or velocity (Figure S13d; $F_{(2,23)}$=0.4404, p=0.6491, $\eta^2$=0.0369). Moreover, center frequency was not different between groups (Figure S13e; $F_{(2,23)}$=1.782, p=0.1908, $\eta^2$=0.1341), nor was center duration (Figure S13f; $F_{(2,23)}$=2.286, p=0.1243, $\eta^2$=0.1658). Wall Frequency (Figure S13g; $F_{(2,23)}$=1.097, p=0.3506, $\eta^2$=0.0871) and duration (Figure S13h; $F_{(2,23)}$=1.903, p=0.1719, $\eta^2$=0.1420) were not different between groups. Moreover, latency to wall was not significantly different between groups (Figure S13i; $F_{(2,23)}$=1.075, p=0.3578, $\eta^2$=0.08549). Overall, these results indicate that overexpressing CRF2 in the BLA does not alter anxiety-like behaviors or locomotion.

Next, to confirm CRF2 was overexpressed by the virus, qPCR was conducted on half the mice. In the BLA (Figure S14f), there was significant effect of probe ($F_{(1,30)}$=15.052, p=0.001, $\eta^2$=0.334), treatment ($F_{(3,30)}$=5.107, p=0.006, $\eta^2$=0.338), and treatment by probe interaction ($F_{(3,30)}$=5.178, p=0.005, $\eta^2$=0.341). Subsequent ANOVA revealed no differences between CRF1 ($F_{(3,15)}$=0.015, p=0.123, $\eta^2$=0.945), but there was between CRF2 ($F_{(3,15)}$=5.157, p=0.012, $\eta^2$=0.508). Post hoc tests revealed that control + veh is significantly different than CRF2 overexpression + Veh (p=0.045) but not CRF2 overexpression + Blebb (p=0.124) or control + blebb (p=1.000). Control + blebb is different than CRF2 overexpression + veh (p=0.032), but not CRF2 overexpression + blebb (p=0.094), and CRF2 overexpression + veh is not different than CRF2 overexpression + blebb (p-0.934). Subsequent ANOVAs also revealed that CRF1 and CRF2 were not different between control + veh ($F_{(1,6)}$=0.014, p=0.911, $\eta^2$=0.002) or control + blebb ($F_{(1,8)}$=0.003, p=0.957, $\eta^2$=0.0003), but was different between CRF2 overexpression + veh ($F_{(1,8)}$=7.305, p=0.027, $\eta^2$=0.477) and CRF2 overexpression + blebb ($F_{(1,8)}$=12.074, p=0.008, $\eta^2$=0.601). In the hypothalamus (Figure S14g), there were no significant differences in probe ($F_{(1,20)}$=1.044, p=0.319, $\eta^2$=0.050), treatment ($F_{(3,20)}$=2.087, p=0.134, $\eta^2$=0.238), or probe by treatment interaction ($F_{(3,20)}$=1.352, p=0.286, $\eta^2$=0.169). The other half of mice were used for placement checks (Figure S15a). These results indicate that the overexpression virus did increase CRF2, but not CRF1, expression in the BLA specifically.

**Figure S1. Mini-pump programmed to mimic METH's clearance curve with COC.** Table with post hoc analysis demonstrating p-values for a) COC's and b) METH's half-life and clearance rates. c) COC, METH, and COC Pump plasma clearance rates (uM).

**Figure S2. Top ten average normalized genes per brain region.** Highly expressed genes in the a-c) BLA, d-f) dHPC and g-i) NAc in mice treated with saline, METH, or COC.

**Figure S3. Relevant DEGs in the BLA following METH compared to COC CPP.** a) Top ten differently expressed genes (DEGs) found per treatment group by brain region comparisons. b) NMII-related transcript changes by brain region in mice treated with METH compared to COC. c) Neuromodulator receptor transcript changes by brain region in mice treated with METH versus COC.

**Figure S4.** CRF2 pathway analyses. a) IPA analysis of CRF signaling and related pathway, and b) its signaling pathway.

**Figure S5. BLA placements.** a) Placements for METH- and COC-treated mice (vehicle: grey, AS2B: black). b) Placements groups (vehicle: grey, AS2B: black). c) Placements for groups (vehicle: grey; AS2B: black). d) Placements for groups (vehicle + blebb: light green; AS2B + blebb: teal).

**Figure S6. METH-associated learning may not require CRF2**. METH CPP following intra-BLA a) vehicle (n=14; $t_{(13)}$=1.612, p=0.1310, $r^2$=0.1666) or b) AS2B (n=14; $t_{(13)}$=1.923, p=0.0766, $r^2$=0.2215). Error bars represent ± SEM.

**Figure S7. Intra-BLA CRF2 does not impact elevated plus maze performance.** a) Placements (no infusion: light grey, vehicle: grey, AS2B: black) and b) methods outline. Corticosterone levels were not different between groups when measured c) ~25 minutes (n = 10-11 per group) or d) ~15 minutes (n = 7-8 per group) after intra-BLA microinfusion. Latency to e) open and f) closed arms. i) Distance moved in EPM was not different between groups, nor was j) frequency in open or k) closed arm or duration in l) open arm or m) closed arm. (n = 9-11 per group). *p<0.05. Error bars represent ± SEM.

**Figure S8 Intra-BLA microinfusions can alter some anxiety-like behaviors in the open field.** a) center frequency, b) center duration, c) wall frequency, d) wall duration, e) latency to center, f) latency to wall, g) total distance traveled, h) total velocity, i) first minute of the session (to examine initial anxiety-like behavior) distance traveled, and j) first minute velocity during test 1. (n=10-11 per group) *p<0.05. Error bars represent ± SEM.

**Figure S9 Microinfusing AS2B into the BLA does little to anxiety-like behaviors in the open field.** (a) center frequency, b) center duration, c) wall frequency, d) wall duration, e) middle frequency, f) middle duration, latency to g) middle, h) center, or i) wall, j) total distance traveled, k) total velocity, l) first minute of distance traveled, and m) first minute of velocity. (n=10-11 per group). *p<0.05, **p<0.01. Error bars represent ± SEM.

**Figure S10. BLA CRF2 agonism did not render a COC-associated memory susceptible to NMII inhibition.** a) Placements (vehicle + vehicle: grey; vehicle + Blebb: green; UCN3 + vehicle: light purple; UCN3 + Blebb: dark purple) and b) methods outline for COC CPP following c) vehicle + vehicle, d) vehicle + blebb, e) UCN3 + vehicle, or f) UCN3 + Blebb (n = 14 per group). *p<0.05, **p<0.01. Error bars represent ± SEM.

**Figure S11. Determining CRF2 overexpression virus optimal parameters**. a) Methods outline. b) CRF1 and CRF2 mRNA levels with different dilutions of AAV5-mCrhr2-alpha in the BLA (all groups, n=5) and c) hypothalamus (n = 4-5). d) Representative images of different infusion dilutions. *p<0.05. Error bars represent ± SEM.

**Figure S12**. **Representative images from optimal viral parameters.** CRF1 and CRF2 mRNA levels with different intra-BLA infusion volumes of AAV5-mCrhr2-alpha in the a) BLA and b) hypothalamus (n = 3-4 per group) c) Representative images of different infusion volumes. *p<0.05. Error bars represent ± SEM.

**Figure S13. There are no changes to anxiety-like behavior when CRF2 is overexpressed in the BLA.** a) total distance traveled and b) velocity did not differ between groups. When the first minute of the session was analyzed, no differences in groups were found in c) distance or d) velocity. e) Center frequency, f) center duration, g) wall frequency, and h) wall duration did not differ between groups in the OF. Neither did i) latency to wall. g). (n = 8-9 per group). Error bars represent ± SEM.

**Figure S14. Overexpressing CRF2 in the BLA did not render a COC-associated memory susceptible to disruption by NMII inhibition.** a) Methods outline. b) Control + vehicle (n=10; $t_{(9)}$=3.024, p=0.0144, $r^2$=0.5040), c) Control + Blebb, (n=10; $t_{(9)}$=3.687, p=0.0049, $r^2$=0.6029), d) CRF2 OE + vehicle (n=10; $t_{(9)}$=3.609, p=0.0057, $r^2$=0.5914), and e) CRF2 OE + Blebb (n=10; $t_{(9)}$ =3.670, p=0.0052, $r^2$=0.5994) all expressed a significant COC-associated memory. qPCR verified that CRF2 OE did result in an increase of CRF2 in the f) BLA but not g) hypothalamus. *p<0.05. **p<0.01. Error bars represent ± SEM.

**Figure S15. Representative images** from a) groups expressing CRF2 overexpression or control virus treated with vehicle or blebb.

Figure S1

a

| COC (p-values) | | | | | | |
|---|---|---|---|---|---|---|
| | 15 minutes | 30 minutes | 1 hour | 2 hours | 4 hours | 8 hours |
| 15 minutes | | 0.000 | 0.000 | 0.000 | 0.000 | 0.000 |
| 30 minutes | 0.000 | | 0.104 | 0.010 | 0.004 | 0.003 |
| 1 hour | 0.000 | 0.104 | | 0.859 | 0.696 | 0.546 |
| 2 hours | 0.000 | 0.010 | 0.859 | | 1.000 | 0.992 |
| 4 hours | 0.000 | 0.004 | 0.696 | 1.000 | | 0.999 |
| 8 hours | 0.000 | 0.003 | 0.546 | 0.992 | 0.999 | |

b

| METH (p-values) | | | | | | | |
|---|---|---|---|---|---|---|---|
| | 15 minutes | 30 minutes | 1 hour | 2 hours | 4 hours | 8 hours | 12 hours |
| 15 minutes | | 0.922 | 0.019 | 0.000 | 0.000 | 0.000 | 0.001 |
| 30 minutes | 0922 | | 0.164 | 0.003 | 0.001 | 0.001 | 0.004 |
| 1 hour | 0.019 | 0.164 | | 0.344 | 0.111 | 0.109 | 0.191 |
| 2 hours | 0.000 | 0.003 | 0.344 | | 0.990 | 0.970 | 0.982 |
| 4 hours | 0.000 | 0.001 | 0.111 | 0.990 | | 1.000 | 1.000 |
| 8 hours | 0.000 | 0.001 | 0.109 | 0.970 | 1.000 | | 1.000 |
| 12 hours | 0.001 | 0.004 | 0.191 | 0.982 | 1.000 | 1.000 | |

c

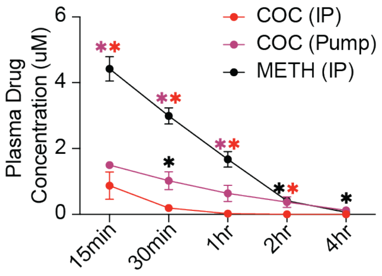

Figure S2

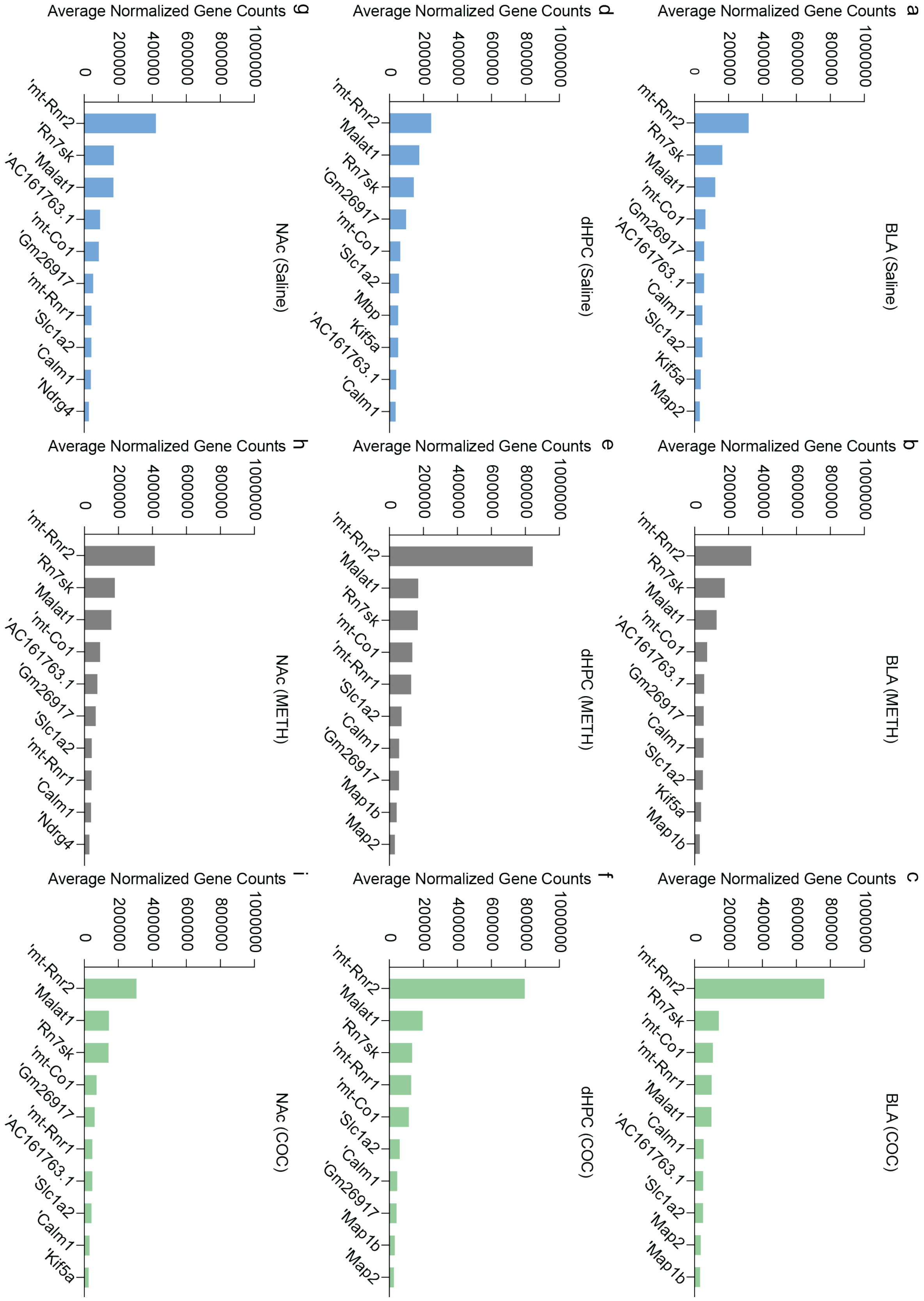
a
BLA (Saline)
b
BLA (METH)
c
BLA (COC)
d
dHPC (Saline)
e
dHPC (METH)
f
dHPC (COC)
g
NAc (Saline)
h
NAc (METH)
i
NAc (COC)
Average Normalized Gene Counts
0
200000
400000
600000
800000
1000000
'mt-Rnr2
'Rn7sk
'Malat1
'mt-Co1
'Gm26917
'AC161763.1
'Calm1
'Slc1a2
'Kif5a
'Map2
'Map1b
'mt-Rnr1
'Mbp
'Ndrg4

### Figure S3

a

| Saline (BLA-NAc) | Saline (BLA-dHPC) | Saline (NAc-dHPC) | METH (BLA-NAc) | METH (BLA-NAc) | METH (NAc-dHPC) | COC (BLA-NAc) | COC (BLA-dHPC) | COC (NAc-dHPC) |
|---|---|---|---|---|---|---|---|---|
| ‘Tnc | ‘Gm2115 | ‘Gm2115 | ‘Chchd4 | ‘Gm26695 | ‘Ak6 | ‘Cyp26b1 | ‘Arhgap6 | ‘Gpr88 |
| ‘Scml4 | ‘Sipa1l3 | ‘Sipa1l3 | ‘Tpbg | ‘Mroh7 | ‘AC102847.1 | ‘Nr2f2 | ‘Gpr88 | ‘Rxrg |
| ‘Crym | ‘Otof | ‘Prlar2b | ‘E030044B06Rik | ‘Trmu | ‘2310011J03Rik | ‘Pcp4l1 | ‘Rxrg | ‘Dnah9 |
| ‘Evc2 | ‘Gap43 | ‘Cpne5 | ‘Gm42449 | ‘Tfb2m | ‘Vamp3 | ‘Serpina9 | ‘Pbx3 | ‘Penk |
| ‘Hgf | ‘Rxrg | ‘Rarb | ‘Tra2b | ‘Fam53b | ‘Zfp7 | ‘Rgs9 | ‘Ankfn1 | ‘Rgs9 |
| ‘Slc6a7 | ‘Oprk1 | ‘Gnal | ‘Ifnar1 | ‘Exosc2 | ‘Ogn | ‘C1ql3 | ‘Penk | ‘Pbx3 |
| ‘Zbtb20 | ‘Rps6ka2 | ‘Oprk1 | ‘Emsy | ‘Liph | ‘Dctn1 | ‘Ankfn1 | ‘Gm2115 | ‘Arhgap6 |
| ‘Cacna1g | ‘Gpr161 | ‘Gpr161 | ‘Grik3 | ‘Gm43267 | ‘Stx7 | ‘Dnah9 | ‘Olfml2b | ‘Tac1 |
| ‘Ak4 | ‘Epha6 | ‘Meis2 | ‘Ilf3 | ‘Gm43267 | ‘Slc26a5 | ‘Gm11639 | ‘4921539H07Rik | ‘Drd2 |
| ‘Rarb | ‘Prkar2b | ‘Epha6 | ‘Ralb | ‘Gm17322 | ‘4930525G20Rik | ‘Frem1 | Unc13c | ‘4921539H07Rik |

b

**NMII-related transcript changes by brain region in mice treated with METH compared to COC**

| Protein | Gene | BLA | | dHPC | | NAc | |
|---|---|---|---|---|---|---|---|
| **RLC's:** | | **pvalue** | **Fold Change** | **pvalue** | **Fold Change** | **pvalue** | **Fold Change** |
| MLC3 | ‘Myl6 | 0.080 | 0.855 | 0.263 | 0.820 | 0.287 | 1.085 |
| MLC2B | ‘Myl12a | 0.005 | 0.723 | 0.117 | 0.722 | 0.861 | 0.979 |
| MLC2 | ‘Myl9 | 0.014 | 0.635 | 0.147 | 0.567 | 0.358 | 0.846 |
| MLC1sa | ‘Myl4 | 0.000 | 0.462 | 0.618 | 0.380 | 0.306 | 1.446 |
| **MLCP subunits:** | | | | | | | |
| PP1 | ‘Ppp1cc | 0.070 | 0.857 | 0.623 | 1.072 | 0.857 | 1.017 |
| **RLC kinases:** | | | | | | | |
| MRCK | ‘Cdc42bpg | 0.213 | 0.700 | 0.214 | 0.613 | 0.175 | 0.821 |
| MRCK | ‘Cdc42bpb | 0.072 | 0.751 | 0.849 | 0.946 | 0.891 | 1.007 |
| MRCK | ‘Cdc42bpa | 0.041 | 1.182 | 0.860 | 1.023 | 0.660 | 1.034 |
| ROCK | ‘Rock2 | 0.093 | 1.256 | 0.568 | 1.135 | 0.877 | 0.976 |
| MLCK | ‘Mylk | 0.638 | 1.049 | 0.008 | 3.885 | 0.905 | 0.842 |
| **MLCP kinases:** | | | | | | | |
| PKG2 | ‘Prkg2 | 0.003 | 1.484 | 0.996 | 1.001 | 0.705 | 0.955 |
| **MLCP inactivators:** | | | | | | | |
| Darpp-32 | ‘Ppp1r1b | 0.019 | 0.574 | 0.793 | 0.938 | 0.617 | 0.927 |
| ROCK | ‘Rock2 | 0.093 | 1.256 | 0.568 | 1.135 | 0.877 | 0.976 |

c

**Neuromodulator receptor transcript changes by brain region in mice treated with METH versus COC**

| | BLA | | dHPC | | NAc | |
|---|---|---|---|---|---|---|
| **Gene** | **pvalue** | **Fold Change** | **pvalue** | **Fold Change** | **pvalue** | **Fold Change** |
| ‘Drd2 | 0.025 | 0.568 | 0.176 | 0.659 | 0.611 | 0.922 |
| ‘Crhr2 | 0.019 | 0.531 | 0.894 | 0.931 | 0.943 | 1.025 |
| ‘Adra1a | 0.029 | 1.313 | 0.739 | 0.918 | 0.654 | 1.057 |
| ‘Adra2b | 0.075 | 0.591 | 0.721 | 1.353 | 0.893 | 0.971 |
| ‘Adra1b | 0.020 | 0.637 | 0.218 | 1.714 | 0.194 | 1.420 |
| ‘Adra2c | 0.006 | 0.607 | 0.866 | 0.943 | 0.104 | 0.816 |
| ‘Adra1d | 0.036 | 0.530 | 0.556 | 0.760 | 0.404 | 1.302 |

Figure S4

a

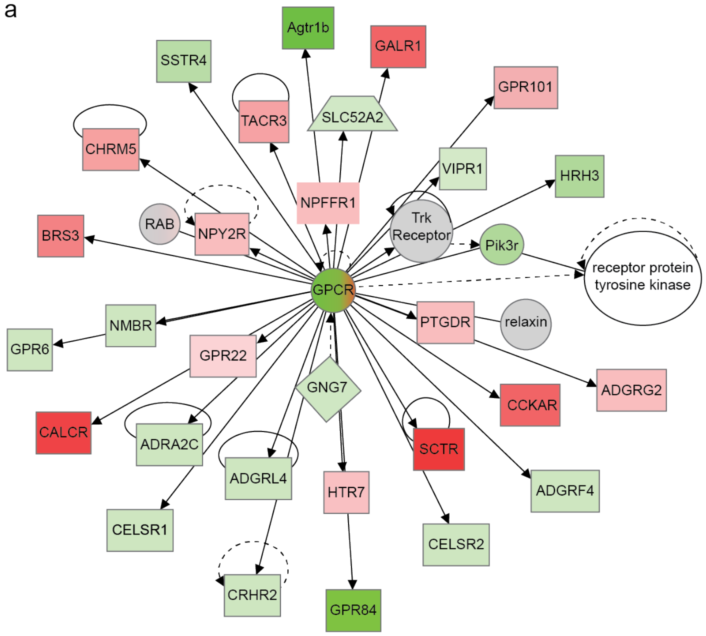
Agtr1b
GALR1
SSTR4
GPR101
TACR3
SLC52A2
CHRM5
VIPR1
HRH3
NPFFR1
RAB
Trk Receptor
BRS3
NPY2R
Pik3r
receptor protein tyrosine kinase
GPCR
NMBR
PTGDR
relaxin
GPR6
GPR22
ADGRG2
GNG7
CCKAR
CALCR
ADRA2C
SCTR
ADGRL4
HTR7
ADGRF4
CELSR1
CELSR2
CRHR2
GPR84

b

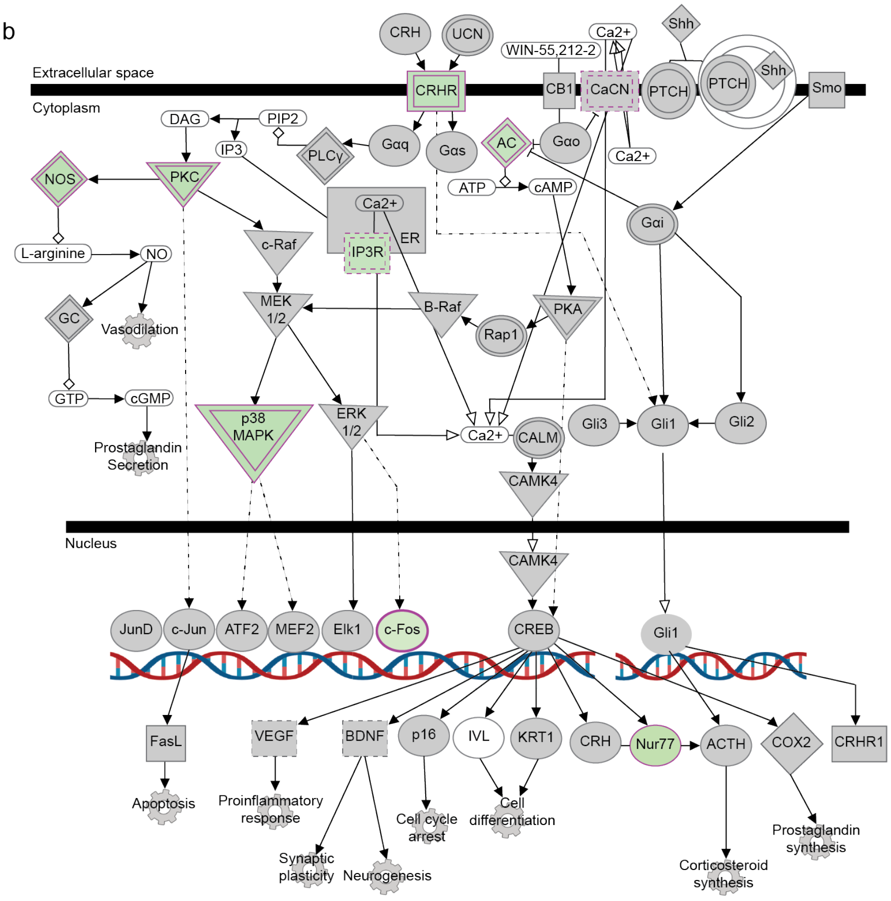
CRH
UCN
Ca2+
Shh
WIN-55,212-2
Extracellular space
CRHR
CB1
CaCN
PTCH
PTCH
Shh
Smo
Cytoplasm
DAG
PIP2
IP3
PLCγ
Gαq
Gαs
AC
Gαo
Ca2+
NOS
PKC
ATP
cAMP
Ca2+
ER
IP3R
Gαi
L-arginine
NO
c-Raf
GC
Vasodilation
MEK 1/2
B-Raf
PKA
Rap1
GTP
cGMP
p38 MAPK
ERK 1/2
Ca2+
CALM
Gli3
Gli1
Gli2
Prostaglandin Secretion
CAMK4
Nucleus
CAMK4
JunD
c-Jun
ATF2
MEF2
Elk1
c-Fos
CREB
Gli1
FasL
VEGF
BDNF
p16
IVL
KRT1
CRH
Nur77
ACTH
COX2
CRHR1
Apoptosis
Proinflammatory response
Cell cycle arrest
Cell differentiation
Prostaglandin synthesis
Synaptic plasticity
Neurogenesis
Corticosteroid synthesis

Figure S5

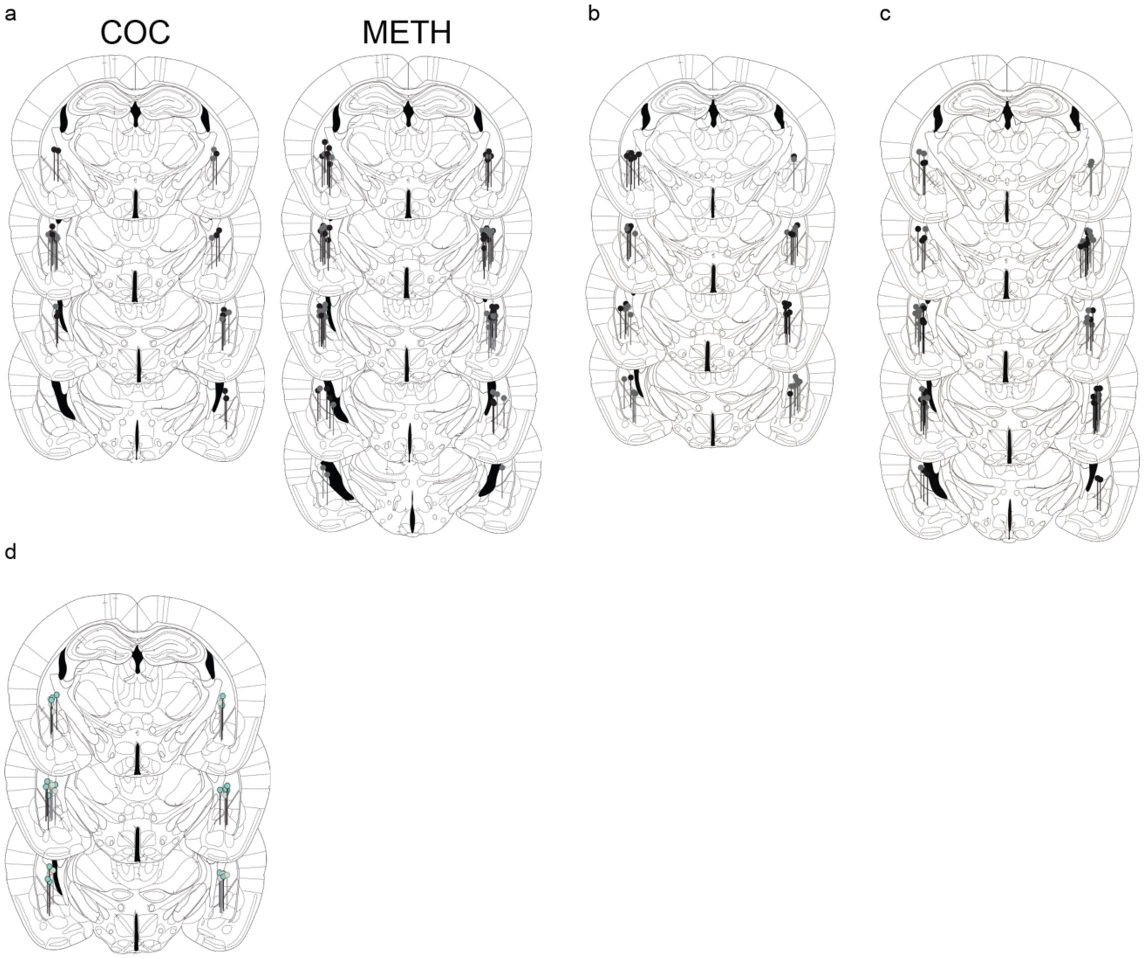

Figure S6

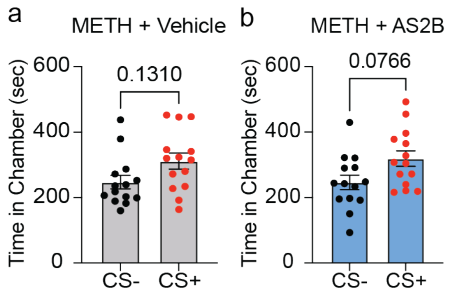
a
METH + Vehicle
0.1310
Time in Chamber (sec)
600
400
200
0
CS-
CS+
b
METH + AS2B
0.0766
Time in Chamber (sec)
600
400
200
0
CS-
CS+

Figure S7

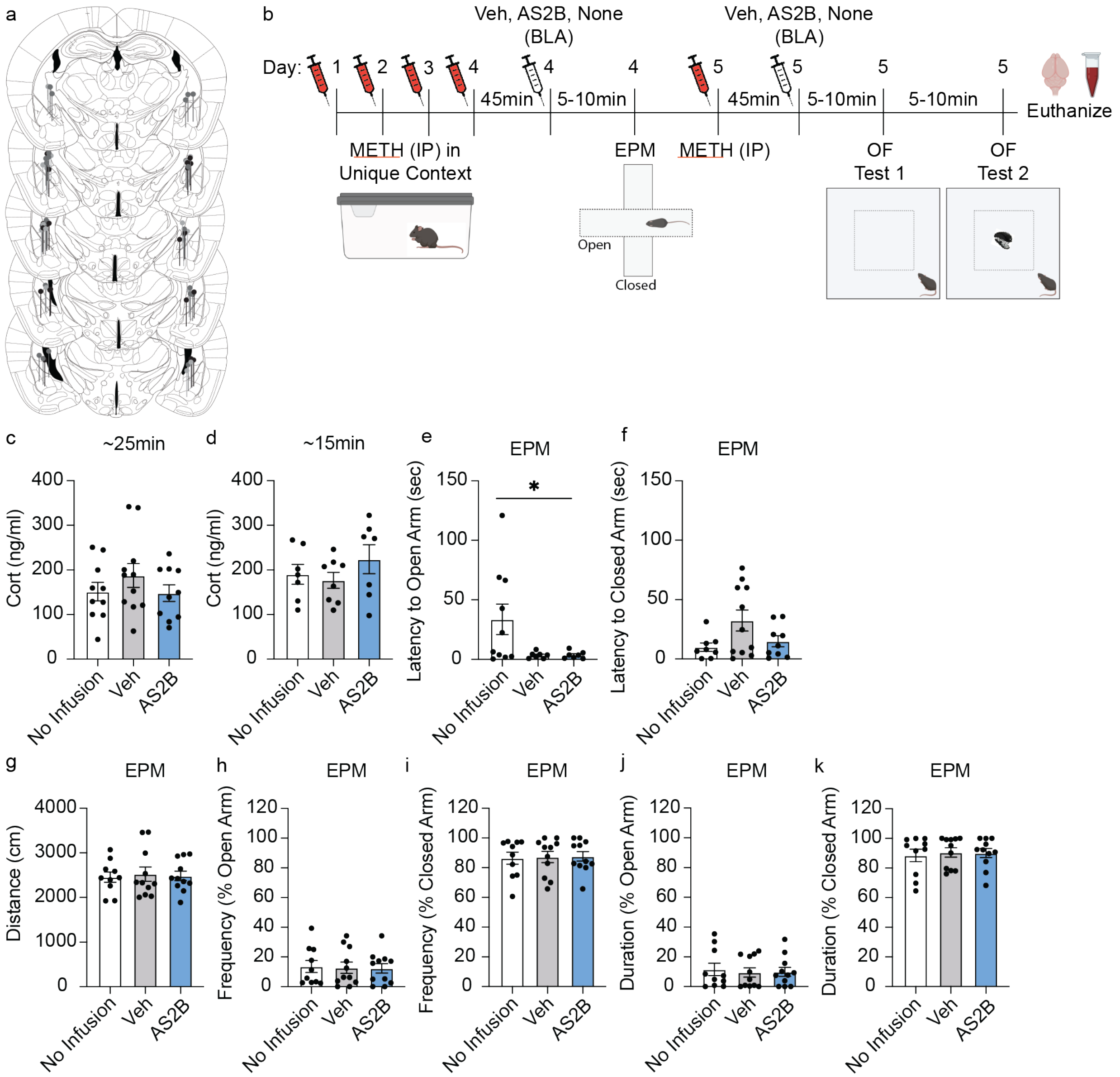
a
b
Veh, AS2B, None (BLA)
Veh, AS2B, None (BLA)
Day: 1 2 3 4 4 4 5 5 5 5
45min
5-10min
45min
5-10min
5-10min
Euthanize
METH (IP) in Unique Context
EPM
METH (IP)
OF Test 1
OF Test 2
Open
Closed
c
~25min
Cort (ng/ml)
No Infusion
Veh
AS2B
d
~15min
Cort (ng/ml)
e
EPM
Latency to Open Arm (sec)
*
f
EPM
Latency to Closed Arm (sec)
g
EPM
Distance (cm)
h
EPM
Frequency (% Open Arm)
i
EPM
Frequency (% Closed Arm)
j
EPM
Duration (% Open Arm)
k
EPM
Duration (% Closed Arm)

Figure S8

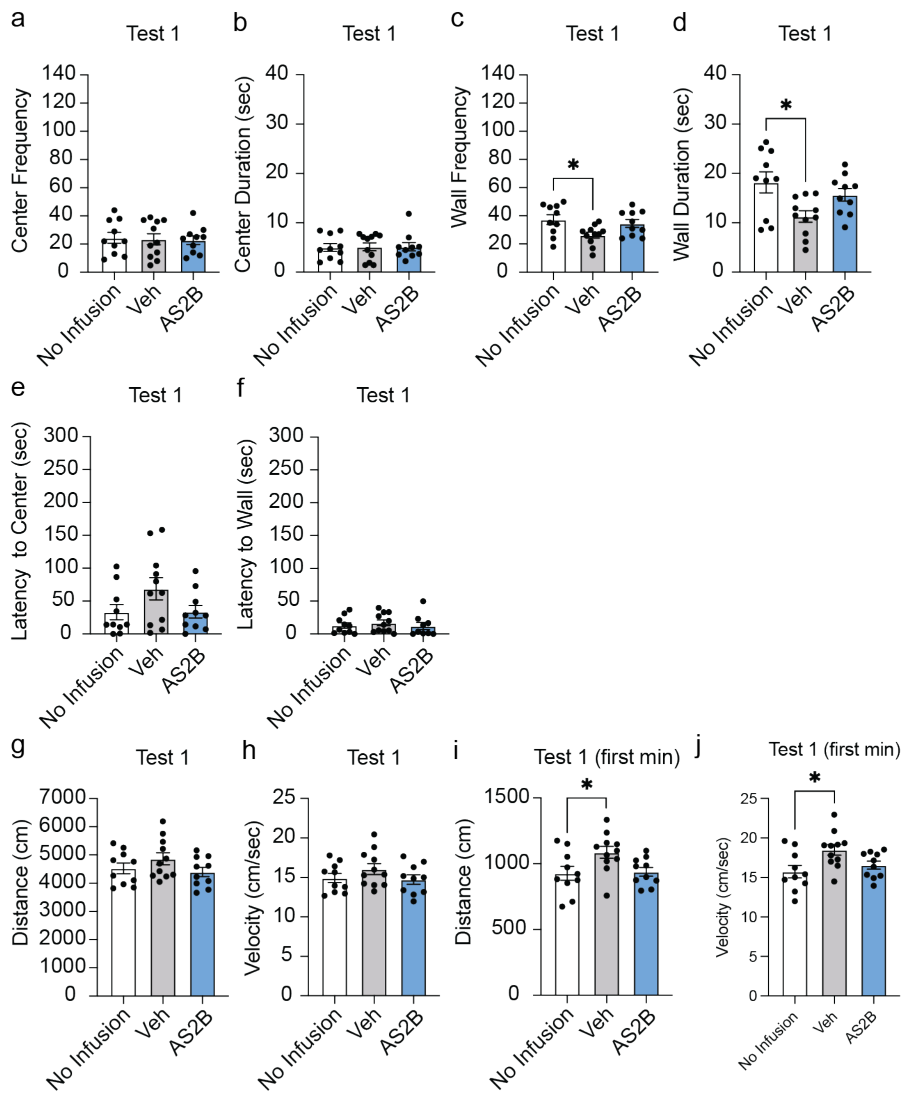
a
Test 1
Center Frequency
No Infusion
Veh
AS2B
b
Test 1
Center Duration (sec)
c
Test 1
Wall Frequency
*
d
Test 1
Wall Duration (sec)
*
e
Test 1
Latency to Center (sec)
f
Test 1
Latency to Wall (sec)
g
Test 1
Distance (cm)
h
Test 1
Velocity (cm/sec)
i
Test 1 (first min)
Distance (cm)
*
j
Test 1 (first min)
Velocity (cm/sec)
*

Figure S9

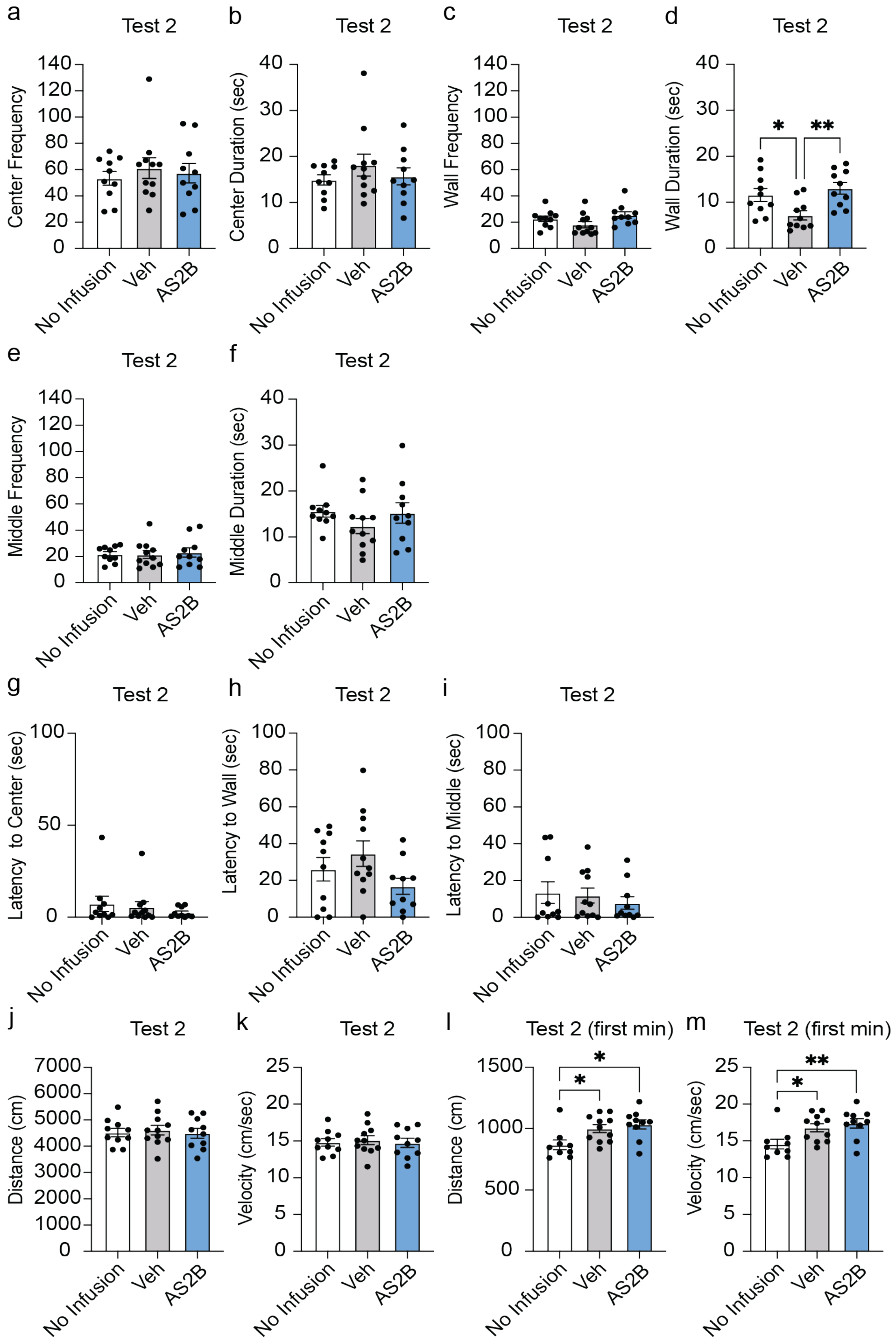
a
Test 2
Center Frequency
140
120
100
80
60
40
20
0
No Infusion
Veh
AS2B
b
Test 2
Center Duration (sec)
40
30
20
10
0
c
Test 2
Wall Frequency
d
Test 2
Wall Duration (sec)
*
**
e
Test 2
Middle Frequency
f
Test 2
Middle Duration (sec)
g
Test 2
Latency to Center (sec)
100
50
0
h
Test 2
Latency to Wall (sec)
i
Test 2
Latency to Middle (sec)
j
Test 2
Distance (cm)
7000
6000
5000
4000
3000
2000
1000
0
k
Test 2
Velocity (cm/sec)
25
20
15
10
5
0
l
Test 2 (first min)
Distance (cm)
1500
1000
500
0
*
*
m
Test 2 (first min)
Velocity (cm/sec)
*
**

Figure S10

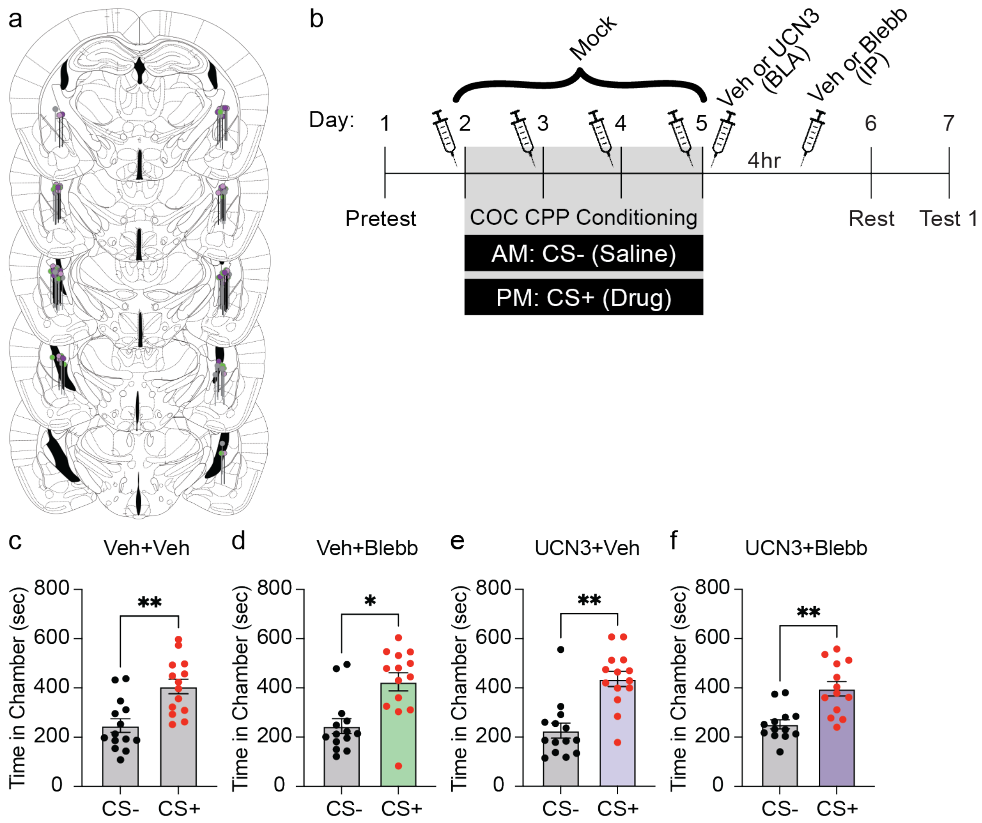
a
b
Mock
Veh or UCN3 (BLA)
Veh or Blebb (IP)
Day: 1 2 3 4 5 6 7
4hr
Pretest
COC CPP Conditioning
AM: CS- (Saline)
PM: CS+ (Drug)
Rest
Test 1
c
Veh+Veh
d
Veh+Blebb
e
UCN3+Veh
f
UCN3+Blebb
Time in Chamber (sec)
0
200
400
600
800
**
*
CS-
CS+

Figure S11

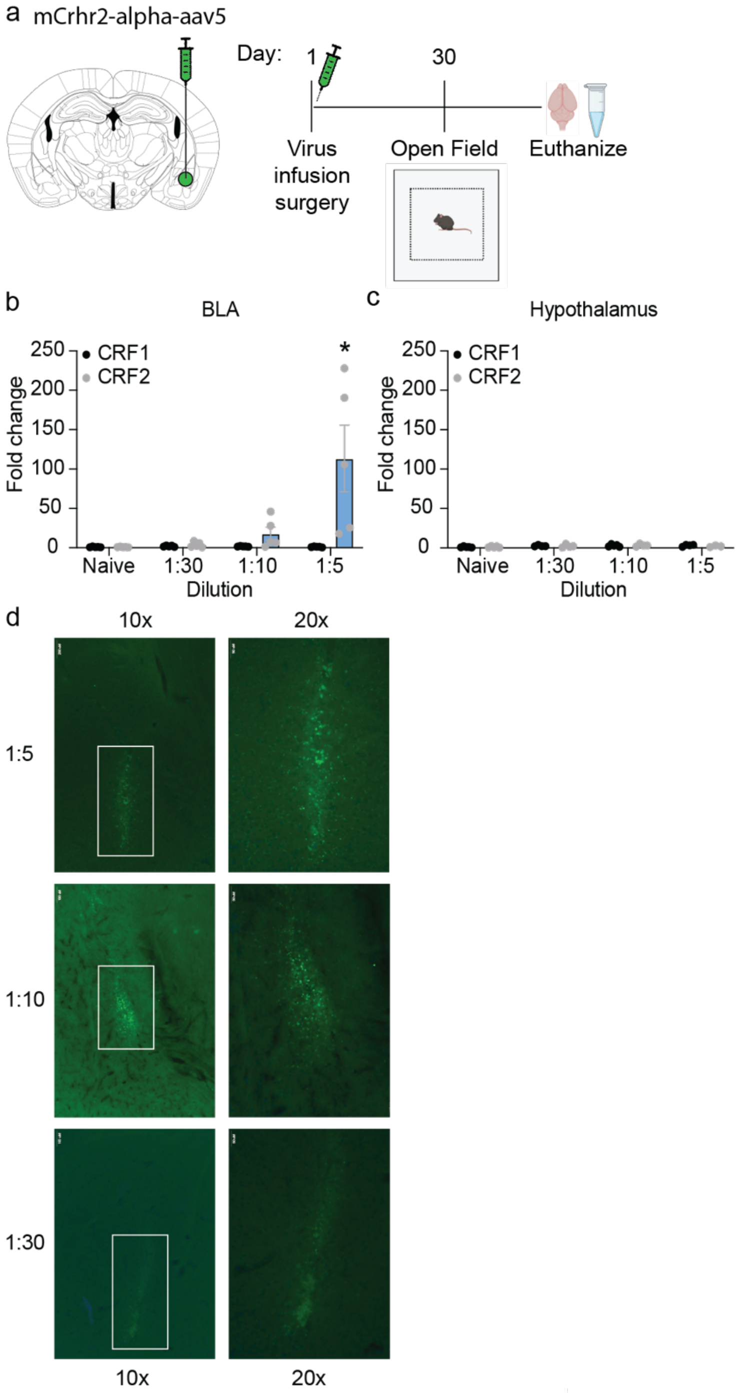
a
mCrhr2-alpha-aav5
Day:
1
30
Virus infusion surgery
Open Field
Euthanize
b
BLA
CRF1
CRF2
Fold change
250
200
150
100
50
0
*
Naive
1:30
1:10
1:5
Dilution
c
Hypothalamus
CRF1
CRF2
Fold change
250
200
150
100
50
0
Naive
1:30
1:10
1:5
Dilution
d
10x
20x
1:5
1:10
1:30
10x
20x

Figure S12

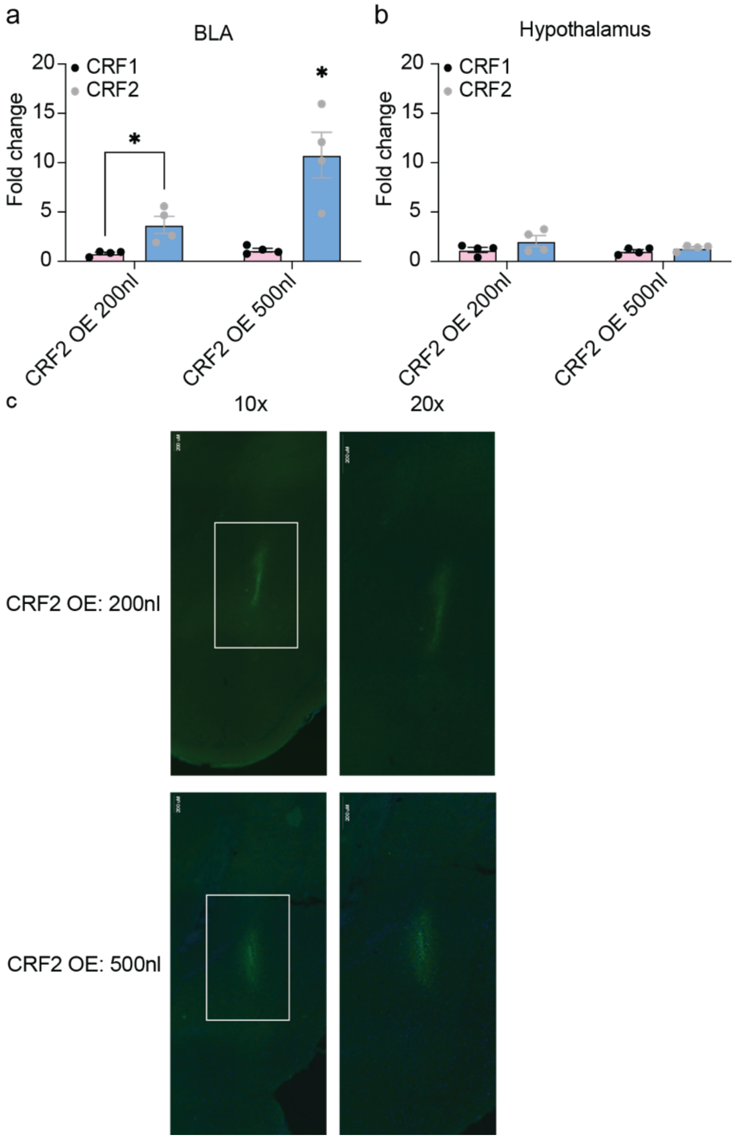
a
BLA
Fold change
20
15
10
5
0
CRF1
CRF2
*
*
CRF2 OE 200nl
CRF2 OE 500nl
b
Hypothalamus
Fold change
20
15
10
5
0
CRF1
CRF2
CRF2 OE 200nl
CRF2 OE 500nl
c
10x
20x
CRF2 OE: 200nl
CRF2 OE: 500nl

Figure S13

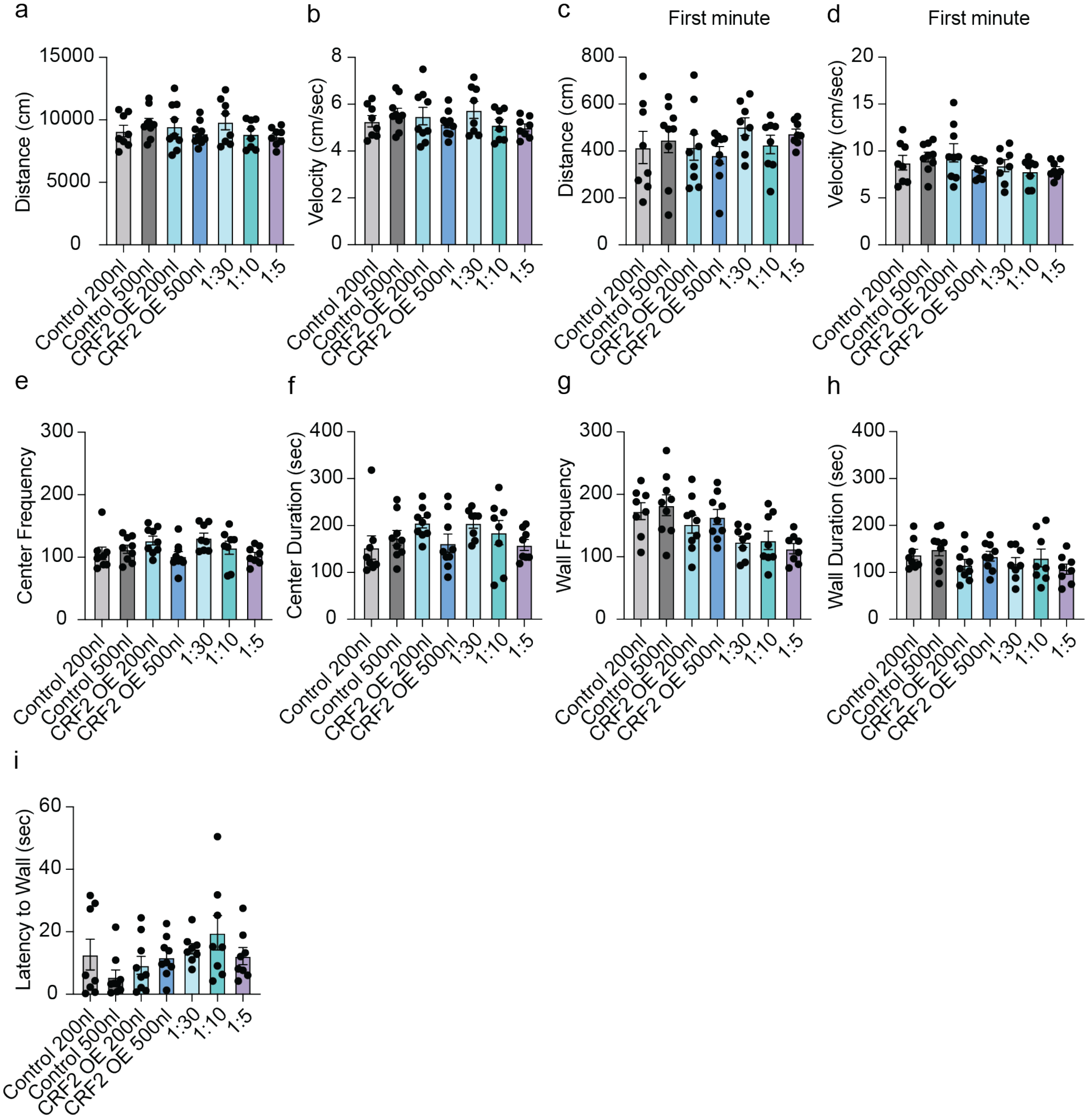
a
Distance (cm)
b
Velocity (cm/sec)
c
First minute
Distance (cm)
d
First minute
Velocity (cm/sec)
e
Center Frequency
f
Center Duration (sec)
g
Wall Frequency
h
Wall Duration (sec)
i
Latency to Wall (sec)
Control 200nl
Control 500nl
CRF2 OE 200nl
CRF2 OE 500nl
1:30
1:10
1:5

Figure S14

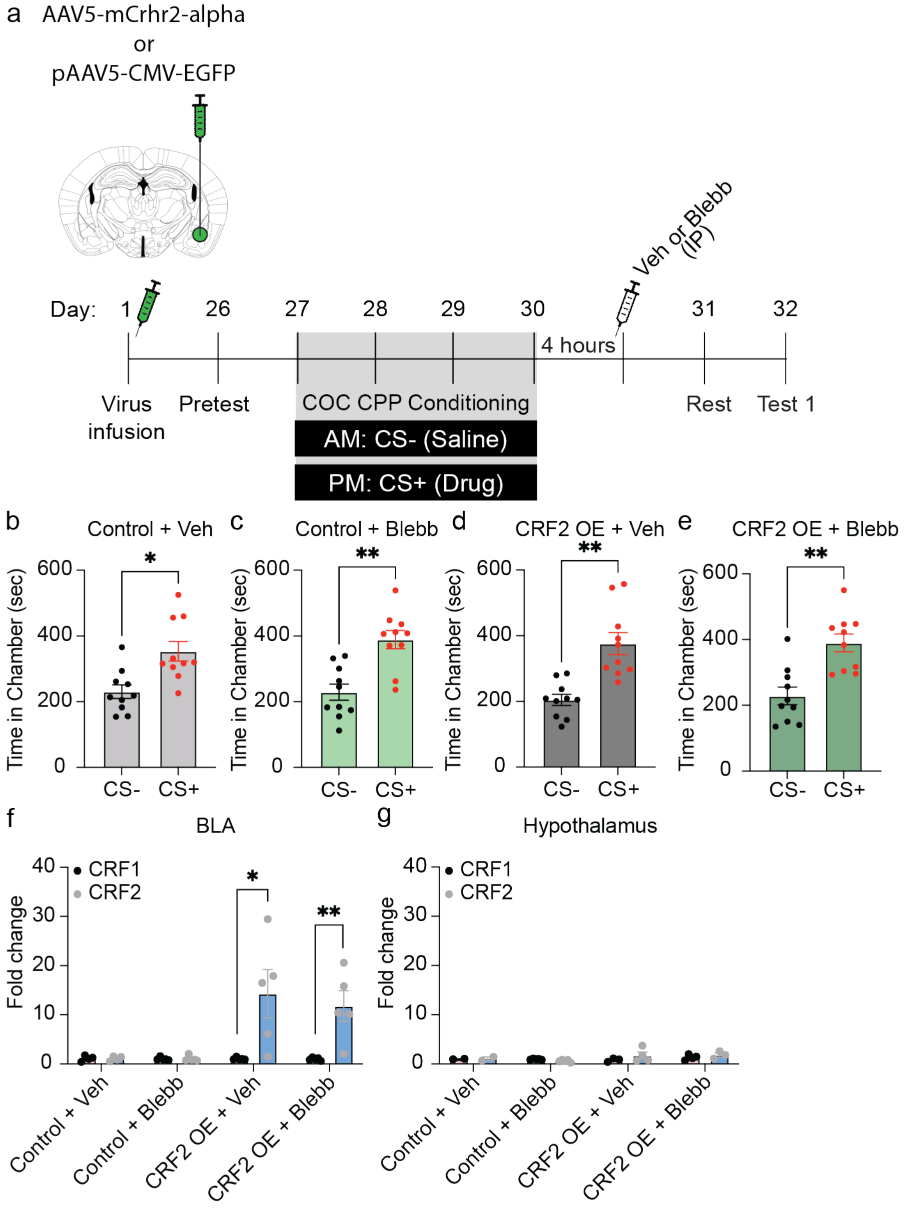
a
AAV5-mCrhr2-alpha
or
pAAV5-CMV-EGFP
Veh or Blebb (IP)
Day: 1 26 27 28 29 30 31 32
4 hours
Virus infusion
Pretest
COC CPP Conditioning
AM: CS- (Saline)
PM: CS+ (Drug)
Rest
Test 1
b
Control + Veh
c
Control + Blebb
d
CRF2 OE + Veh
e
CRF2 OE + Blebb
Time in Chamber (sec)
600
400
200
0
CS-
CS+
*
**
f
BLA
g
Hypothalamus
Fold change
40
30
20
10
0
CRF1
CRF2
Control + Veh
Control + Blebb
CRF2 OE + Veh
CRF2 OE + Blebb

Figure S15

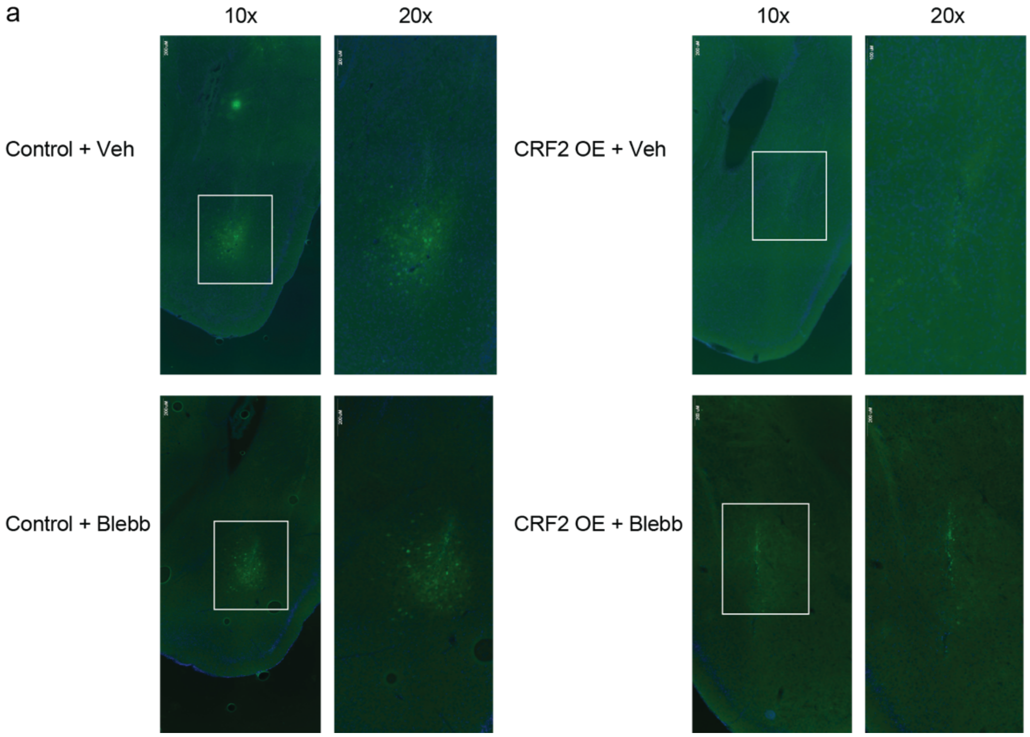
a
10x
20x
10x
20x
Control + Veh
CRF2 OE + Veh
Control + Blebb
CRF2 OE + Blebb
